# A natural variation-based screen in mouse cells reveals USF2 as a regulator of the DNA damage response and cellular senescence

**DOI:** 10.1101/2022.04.21.489100

**Authors:** Taekyu Kang, Emily C. Moore, Emily E. K. Kopania, Christina D. King, Birgit Schilling, Judith Campisi, Jeffrey M. Good, Rachel B. Brem

## Abstract

Cellular senescence is a program of cell cycle arrest, apoptosis resistance, and cytokine release induced by stress exposure in metazoan cells. Landmark studies in laboratory mice have characterized a number of master senescence regulators, including p16^INK4a^, p21, NF-kB, p53, and C/EBPβ. To discover other molecular players in senescence, we developed a screening approach to harness the evolutionary divergence between mouse species. We found that primary cells from the Mediterranean mouse *Mus spretus*, when treated with DNA damage to induce senescence, produced less cytokine and had less-active lysosomes than cells from laboratory *M. musculus*. We used allele-specific expression profiling to catalog senescence-dependent *cis*-regulatory variation between the species at thousands of genes. We then tested for correlation between these expression changes and interspecies sequence variants in the binding sites of transcription factors. Among the emergent candidate senescence regulators, we chose a little-studied cell cycle factor, USF2, for molecular validation. In acute irradiation experiments, cells lacking USF2 had compromised DNA damage repair and response. Longer-term senescent cultures without USF2 mounted an exaggerated senescence regulatory program—shutting down cell cycle and DNA repair pathways, and turning up cytokine expression, more avidly than wild-type. We interpret these findings under a model of pro-repair, anti-senescence regulatory function by USF2. Our study affords new insights into the mechanisms by which cells commit to senescence, and serves as a validated proof of concept for natural variation-based regulator screens.

## Introduction

Metazoan cells of many types, upon exposure to stress, can enter a senescence program, in which they stop dividing, become refractory to apoptosis, and release soluble inflammation and tissue remodeling factors termed the senescence-associated secretory phenotype (SASP) (1– 4). The resulting acute immune response can clear debris, promote wound healing, and/or suppress tumorigenesis (5–8). However, during aging, senescent cells can remain long past any initial triggering event, resulting in chronic inflammation that damages the surrounding tissue (5,9–12). Landmark work has revealed the benefits of eliminating senescent cells to treat age-related pathologies and boost median lifespan (7,8,13,14).

Establishment of the senescent state and the activity of senescent cells hinge in large part on gene regulatory events. Finding molecular players that control this process is an active area of research. Now-classic work has implicated p16^INK4a^ and p21 in the repression of pro-cell cycle genes and promotion of growth arrest (15) after DNA damage, and NF-kB, p53, and C/EBPβ as regulators of the SASP (16). However, given the complexity of the senescence program, many more regulators likely remain to be identified. Indeed, bioinformatic approaches have identified dozens of other transcription factor candidates in senescence (17–24), many of which remain unvalidated (but see (18–21) for recent discoveries of the roles of DLX2, FOXO3 and AP-1).

We set out to develop a novel approach to survey transcription factors that play a role in cellular senescence, with the potential for increased specificity relative to traditional genomic screens. Our strategy took advantage of the natural genetic variation in senescence gene expression, and transcription factor binding sites, across mouse species. Among the top hits from this analysis, we chose the under-studied factor USF2 for validation experiments, focusing on gene regulation and cellular phenotypes during senescence induction and maintenance.

## Methods

### Primary cell extraction and culture

Wild-derived lines of *Mus musculus* (PWK/PhJ), *Mus spretus* (STF/Pas), and their interspecies F1 hybrids (*M. musculus* x *M. spretus*), were maintained in standard conditions under Montana Institutional Animal Care and Use Committee protocol number 062-1JGDBS-120418. For each genotype, five tails from two males and three females aged 3 to 5 months were collected into chilled Dulbecco’s Modified Eagle Medium (DMEM) and shipped to the Buck Institute/UC Berkeley for further processing. *M. domesticus* TUCA, from Tucson, Arizona, in their 40th generation of sib-sib mating, and MANB, from Manaus, Brazil, in their 25th generation of sib-sib mating, were maintained in standard conditions under UC Berkeley Institutional Animal Care and Use Committee protocol number AUP-2016-03-8548-2. For each genotype, two tails from female mice less than 10 weeks old were collected as above. No blinding was required for tail collection. Primary tail fibroblasts were extracted from the cuttings essentially as described (25). For experiments in wild-type *M. musculus*, *M. spretus*, *M. domesticus*, and F1 hybrid cells we considered the culture from each individual animal to represent one biological replicate of the respective genotype.

### Irradiation treatment

To treat a given cell culture replicate with ionizing radiation (3,26,27) for a cell-biological assay or omics profiling, we proceeded as follows. The day before irradiation, cells were seeded at 60-70% confluency and incubated in a 37°C humidified incubator at 3% O_2_ and 10% CO_2_ overnight in complete medium. The next day, a subset of cells was collected and used as input into the respective experiment as the unirradiated control. The remainder of the culture was transferred into an X-RAD 320 X-Ray Biological Irradiator and treated with 15 Gy of X-ray irradiation. Cultures were then placed back into the 37°C humidified incubator at 3% O_2_ and 10% CO_2_ until sampling at 6 hours for marker assays and RNA-seq focused on acute DNA damage response, or 7, 10 or 20 days for marker assays and RNA-seq and proteomics focused on senescence (in which case the medium was replaced 6-8 hours after irradiation, then every 48 hours for the remainder of the experiment), as detailed below.

### Senescence marker assays

For a given replicate culture after irradiation and incubation (see above), senescence-associated β-galactosidase activity was measured using the BioVision Inc. Senescence Detection Kit (cat. #K320): cells were fixed, permeabilized, and incubated with the staining solution containing X-gal overnight. Multiple images were taken the following day using a brightfield microscope, and the image names were randomized before the proportion of β-galactosidase-positive cells was counted manually to remove potential sources of bias. Cultures were considered to be senescent if they showed less than 10% EdU incorporation (see below) and over 90% β-galactosidase-positive cells. In the species comparison of Figure 1 we subjected two technical replicate cultures of each of three biological replicates per purebred species to irradiation (see above) followed by β-galactosidase assays at the indicated timepoints. In Figure 6C, we carried out irradiation and β-galactosidase assays as above at day 7 after irradiation for two technical replicates from each of two biological replicates of *M. musculus* cells infected with lentivirus harboring the scrambled control and two of each *Usf2* knockdown (see below). In Supplemental Figure S1, data from each purebred cell lines were collected on day 10 following irradiation. In Supplemental Figure S4, data for purebreds were from Figure 2 at day 7; separately, for the interspecies F1 hybrid, we carried out irradiation and β-galactosidase assays as above for two technical replicates from one biological replicate.

**Figure 1:**
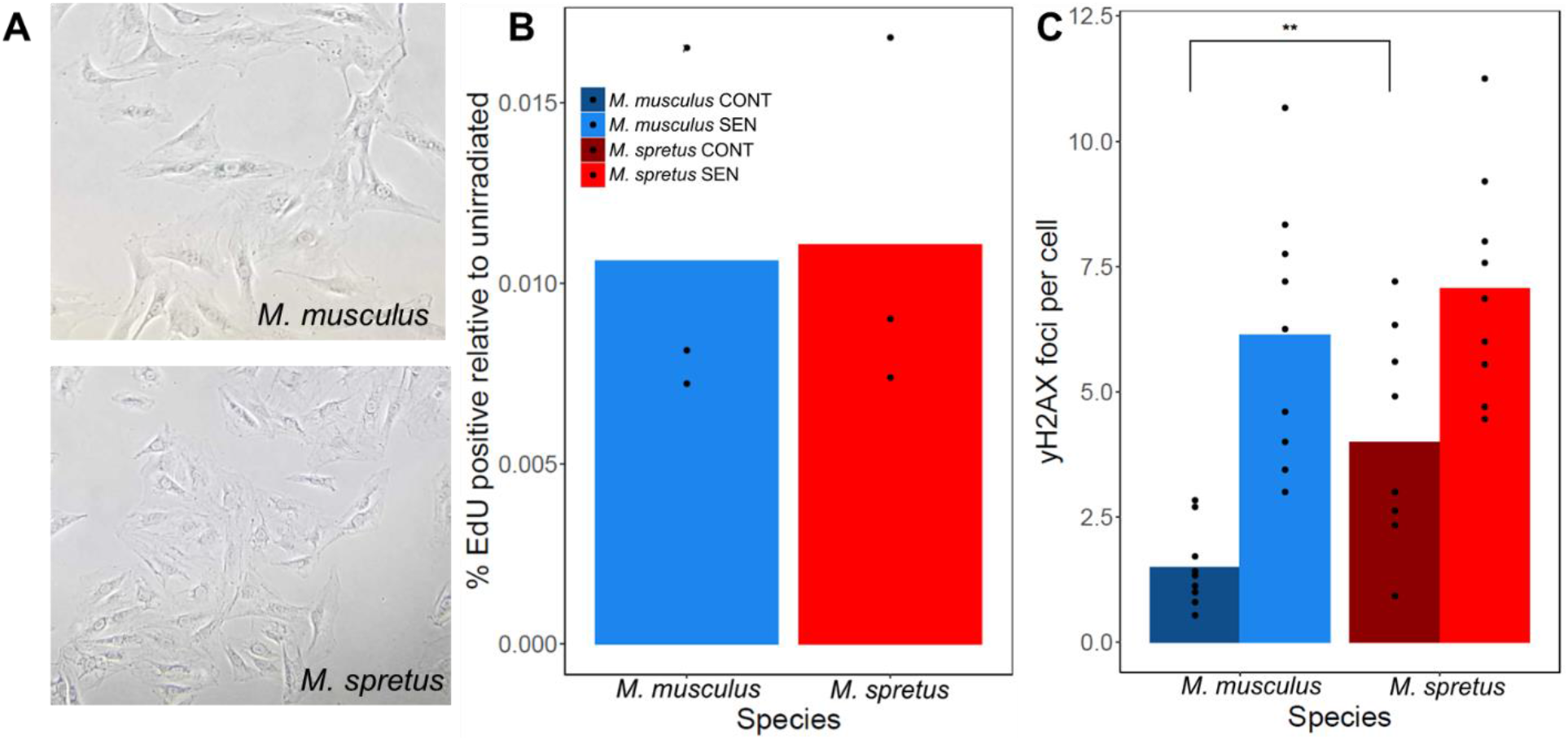
Cells from both *M. musculus* and *M. spretus* mice display expected morphologies of senescence. (A) Representative images of senescent primary fibroblasts from *M. musculus* (above) and *M. spretus* (below) seven days after irradiation exhibiting a flattened and enlarged morphology. (B) Each column reports the average percentage of cells with EdU incorporation seven days after IR treatment (SEN) set relative to the same in unirradiated controls (CONT) for each species reported in (A). For a given column, points report biological and technical replicates (*M. musculus n* = 3, *M. spretus n* = 3). (C) Each columns reports the average number of γH2AX foci per cell for each species as reported in (B). For a given column, points represent biological and technical replicates (*M. musculus n* = 9, *M. spretus n* = 9). **, *p* < 0.01, one-tailed Wilcoxon test comparing species.

**Figure 2:**
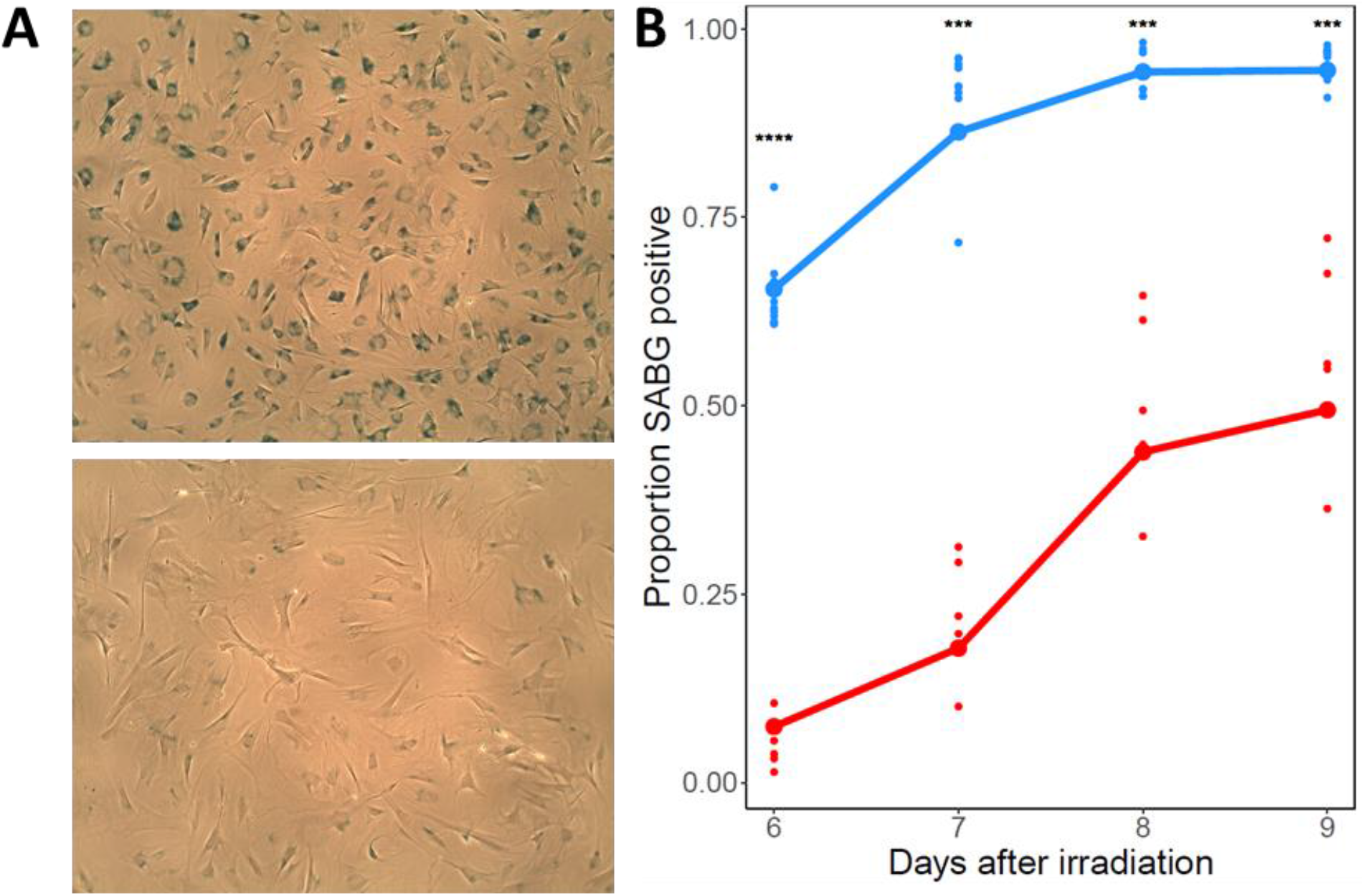
Senescent *M. spretus* cells exhibit lower β-galactosidase activity. (A) Representative images of senescent primary fibroblasts from *M. musculus* (above) and *M. spretus* (below) stained with the β-galactosidase indicator X-Gal seven days after irradiation. (B) Each trace reports results from a timecourse of X-Gal staining assays of primary senescent cells of the indicated species as in (A). The *y*-axis reports the proportion of cells stained positive for senescence-associated β-galactosidase (SABG) activity. In a given column, small points report biological and technical replicates and large points report their average (*M. musculus n* = 9, *M. spretus n* = 5). ***, *p* < 0.001, ****, *p* < 0.0001, one-tailed Wilcoxon test comparing species.

### RNA collection and sequencing

For a given replicate culture of a given genotype, either before irradiation or 6 hours, 10 days, or 20 days after irradiation (see above), cells were treated with TRIzol™ and RNA was extracted using chloroform and ethanol precipitation. The RNA was then further purified using the Qiagen RNeasy Kit (cat. #74004) for DNase treatment and column cleanup. The purity of the extracted RNA was verified using a NanoDrop ND-1000 Spectrophotometer; all samples had 260/280 and 260/230 ratios greater than 2.0. Purified RNA in distilled RNase/DNase free water was snap frozen using dry ice and stored at −80°C. Samples were then either transferred to the QB3 Genomics core at University of California, Berkeley for library prep and sequencing on 150PE NovaSeq S4 or shipped to Novogene Co. (Sacramento, CA) for the same. Both facilities provided 25 million paired end reads per sample. For expression profiles of purebred cells and *M. musculus* x *M. spretus* F1 hybrid cells, we subjected three biological replicates of a given genotype to irradiation (see above) followed by RNA-seq at the indicated timepoints. To assess transcriptional impacts of *Usf2* knockdown we carried out irradiation and RNA isolation as above for two biological replicates of *M. musculus* cells infected with lentivirus harboring the scrambled control and two of each *Usf2* knockdown (see below).

### Pseudogenome and VCF generation

As publicly available annotations for *M. musculus* and *M. spretus* are in the context of their reference genomes (GRCm38.96 and SPRET_EiJ_v1.96 respectively), custom pseudogenomes for strains PWK and STF were generated and used for this study. For the PWK pseudogenome, variant calls between PWK and the reference genome in the form of a variant call file (VCF) were downloaded from the Sanger Mouse Genomes Project database (https://www.sanger.ac.uk/data/mouse-genomes-project/). A pseudogenome using the VCF and the GRCm38.96 reference genome was created using bcftools v1.9 (28). To generate shotgun sequence data for the STF pseudogenome, DNA was extracted from *M. spretus* liver tissue using Qiagen DNeasy spin columns (cat. #69506). The sample was sheared via sonication (Covaris E220), and prepared using the New England Biolabs NEBNext Ultra DNA Library kit (cat. #E7370L). The final library was sequenced on a single lane of 150bp PE Illumina HiSeq X at Novogene, Inc. The latter reads were aligned to the SPRET_EiJ_v1.96 reference genome using bowtie v2.2.3 (29), and a VCF was generated using bfctools mpileup and filtered for quality, depth, and SNPs using vcfutils (30,31). The VCF was then used with the reference genome to create a STF pseudogenome using bcftools. The pseudogenomes were verified by identifying a number of called variants by hand. A VCF of variants between the STF and PWK pseudogenomes was generated as above aligning the STF whole genome sequencing reads to the PWK pseudogenome.

### RNA-seq processing

In transcriptional profiles of *M. musculus* x *M. spretus* F1 hybrid cells, sequencing reads from a given replicate were aligned to a concatenated PWK/STF pseudogenome using tophat v2.1.1 (32) allowing for zero mismatches to ensure allele specific mapping. Alignments were then filtered for uniquely mapped reads using samtools v1.3.1, and gene counts were generated using HTSeq v0.11.2 (33) and genome annotations (GRCm38.96, SPRET_EiJ_v1.96) for both species from Ensembl. Counts were then converted to transcripts per million (TPM) using custom R scripts, and genes were filtered for those showing counts in more than half the samples sequenced.

In transcriptional profiles of purebred *M. musculus* wild-type cells and *M. musculus* cells infected with lentiviruses harboring shRNAs, RNA-seq processing was as above but mapping was to the PWK pseudogenome only; for profiles of purebred *M. spretus* wild-type cells, mapping was to the STF pseudogenome only.

Data for the average TPM across biological replicates for purebred parents and interspecific F1 hybrid are reported in Supplemental Table S1. Average TPM across biological replicates for shRNA-treated *M. musculus* cells are reported in Supplemental Table S6.

### Gene Ontology enrichment analysis of RNA-seq data

For the comparison of transcriptional profiles of *M. musculus* and *M. spretus* purebred cells, for each gene in turn we tabulated the average TPM count from each species across replicates, and then took the log_2_ of the ratio of these averages, r_sen,true_. We downloaded Gene Ontology annotations from the AmiGO 2 (34,35) database and filtered for those with supporting biological data. For each term, we summed the r_sen,true_ values across all genes of the term, yielding s_sen,true_. To assess the enrichment for high or low values of this sum, we first took the absolute value, |s_sen,true_|. We then sampled, from the total set of genes with expression data, a random set of the same number as that in the true data for the term; we calculated the species difference r_sen,rand_ for each such gene and the absolute value of the sum over them all, |s_sen,rand_|. We used as a *p*-value the proportion of 10,000 resampled data sets in which |s_sen,true_| > |s_sen,rand_|.

For analysis of the impact of *Usf2* knockdown on expression before or 6 hours, 10 days, or 20 days after irradiation (see below), Gene Ontology enrichment tests were as above except that we took the ratio, for a given gene, between the average expression in purebred *M. musculus* (PWK) cells infected with lentivirus harboring scrambled shRNA and the analogous quantity across both *Usf2*-targeting shRNA treatments.

### Proteomic analysis of secreted proteins

For a given replicate culture, either before irradiation or 10 days after irradiation (see above), cells were washed three times with PBS and incubated with serum and phenol red free DMEM containing 1% pen-strep for 24 hours. The following day the conditioned medium was collected and passed through a 0.45 μm filter to remove cellular debris. The conditioned medium was placed in a −80°C freezer for storage before use as input into proteomic profiling (see below). For proteomic profiles of purebred cells, we carried out this procedure for three technical replicate cultures of one biological replicate per species.

Sample processing for quantitative proteomic analysis via mass spectrometry was performed as described in (36).

### Transcriptomic screen for senescence regulators

To associate expression variation in genes with sequence variation in their upstream binding sites for a given transcription factor, we proceeded as follows. From RNA-seq profiling of *M. musculus* x *M. spretus* F1 hybrid cells (see above), we used the TPM counts for each parent species’ allele from each replicate profile from control and senescent conditions as input into a two-factor ANOVA. A given gene was categorized as exhibiting senescence-associated differential allele specific expression if the interaction F statistic value from this ANOVA was among the top 25% of all genes tested. Separately, we used compiled data from chromatin immunoprecipitation via high-throughput sequencing from the Gene Transcription Regulation Database (GTRD) (37) to identify all experimentally determined transcription factor (TF) binding sites located within a 5kb window upstream of the transcriptional start site for each gene in turn in the *M. musculus* genome; we refer to the downstream gene of each such binding location as the target of the TF. This calculation used *M. musculus* gene start sites from the Ensembl GRCm38.96 GFF. Next, for each binding site, we used the VCF between PWK and STF pseudogenomes (see above) to identify single nucleotide variants between PWK and STF in the binding site locus. Now, for all the target genes of a given TF, we categorized them as having sequence variants or not in the respective binding site, and exhibiting senescence-associated differential allele-specific expression. We eliminated from further consideration any TF with fewer than 250 target genes in each of the four categories. For all remaining TFs, the 2 × 2 contingency table was used as input into a Fisher’s exact test with Benjamini-Hochberg multiple testing correction.

### *Usf2* shRNA vector design, construction and application

*Usf2* knockdown shRNA sequences were obtained from the Broad Institute Genetic Perturbation Portal (https://portals.broadinstitute.org/gpp/public/). Two shRNA sequences for *Usf2* (CCGGGCAAGACAGGAGCAAGTAAAGCTCGAGCTTTACTTGCTCCTGTCTTGCTTTTTTGAAT; CCGGACAAGGAGACATAATGCATTTCTCGAG-AAATGCATTATGTCTCCTTGTTTTTTTGAAT), and, separately, a scrambled control sequence (CCTAAGGTTAAGTCGCCCTCGCTCGAGCGAGGGCGACTTAACCTTAGG, Addgene cat. #1864), were each cloned into pLKO.1 puro lentiviral vectors (Addgene cat. #8453). Lentiviral particles containing each of the shRNA constructs were generated by calcium phosphate co-transfection of HEK 293T cells with the shRNA pLKO.1 puro vectors and separate pMDLg/pRRE packaging and pCMV-VSV-G envelope plasmids generously provided by Dr. Marius Walter of the Verdin Lab at the Buck Institute. The number of viral particles generated was determined using the Origene One-Wash™ Lentivirus Titer Kit, p24 ELISA (cat. #TR30038). These particles were used to infect two biological replicates of purebred *M. musculus* (PWK) primary tail fibroblasts at a multiplicity of infection of 5 with 4 μg/mL of polybrene, and infected cells were selected by incubating with 2 μg/mL puromycin for 10 days, changing media and antibiotic every other day. Knockdown of *Usf2* was determined by qPCR, using *Usf2* qPCR primer sequences chosen through NCBI Primer Blast, filtering for those spanning an exon-exon junction. The primer pair with the same efficiency (calculated as 10^(−1/slope)^ when plotting log concentration of template cDNA versus Ct) as the internal control *Actb* qPCR primers was chosen: *Usf2* forward 5’ TTCGGCGACCACAATATCCAG 3’, *Usf2* reverse 5’ TTCGGCGACCACAATATCCAG 3’, *Actb* forward 5’ CAACCGTGAAAAGATGACCC 3’, *Actb* reverse 5’ GTAGATGGGCACAGTGTGGG 3’. *Usf2* expression was calculated using the Delta-Delta Ct method (38).

### Cell proliferation and DNA damage assays

For each of two biological replicates of purebred *M. musculus* (PWK) cells infected with lentivirus harboring the scrambled control and two of each *Usf2* knockdown, either before irradiation or 6 hours after irradiation (see above), we measured cell proliferation and DNA damage response as follows.

For a given replicate, DNA synthesis was measured via 5-ethynyl-2′-deoxyuridine (EdU) incorporation assays using the Invitrogen™ Click-iT™ Edu Alexa Fluor™ 488 Flow Cytometry Assay Kit (cat. #C10420). Comet assays were carried out to measure levels of DNA double stranded breaks for a given replicate culture as described (39).

For H2AX assays, for a given replicate, cells were fixed, permeabilized, and blocked, then incubated with 1 μg/mL of primary antibodies specific to phosphorylated (Ser 139) H2AX (cat. # sc-517348, Santa Cruz Biotechnology) in 3% BSA overnight at 4°C. The following day the cells were washed in PBS three times before incubating with 2 μg/mL of Alexa 488 secondary antibodies purchased from Invitrogen (cat. # A11001) for two hours at room temperature. Cells were washed three times with PBS then incubated with 0.5 μg/mL DAPI for 5 minutes at room temperature. The cells were washed once more with PBS before mounting for imaging. Multiple representative confocal images of each sample were taken using a Zeiss LSM 710 AxioObserver, and processed with ImageJ (40).

### Multivariate ANOVA of irradiation and senescence timecourse

To identify genes whose expression changed in wild-type cells through irradiation and senescence, we used RNA-seq profiling data from purebred *M. musculus* (PWK) cells harboring a scrambled shRNA before and 6 hours, 10 days, and 20 days after irradiation (see Supplemental Table S6) as input into a multivariate ANOVA test.

### MERLIN regulatory network reconstruction

To reconstruct a regulatory network we used RNA-seq profiling data for purebred *M. musculus* (PWK) cells harboring a scrambled shRNA or a *Usf2*-targeting RNA, before and 6 hours, 10 days and 20 days after irradiation, as input into MERLIN (41) with default settings. Analysis used a catalog of murine transcription factors from the Gene Transcription Regulation Database (37).

## Results

### High levels of senescence markers in *M. musculus* fibroblasts relative to other mice

To study natural variation in senescence phenotypes, we made use of a classic *in vitro* cell model of senescence, namely primary fibroblasts from mouse tail skin treated with 15 Gy of ionizing radiation (IR) (3,26). We isolated primary tail fibroblasts from the PWK and STF wild-derived purebred lines of *Mus musculus musculus* (hereafter *M. musculus*) and *M. spretus* respectively. Seven days after IR treatment, cells from both species had arrested growth and exhibited the expected flattened and enlarged morphologies of senescent cells (Figure 1A-B). Assaying these cultures for γH2AX foci, a marker of the DNA damage response and a hallmark of long-term senescence (42,43), we found that counts were indistinguishable in primary *M. musculus* and *M. spretus* fibroblasts after irradiation, though higher in the latter in the absence of treatment (Figure 1C). These data indicated that cells of both species had mounted the DNA damage response and entered the senescent state in our treatment and incubation regime.

To begin to compare the senescence program between cells of *M. musculus* and *M. spretus*, we assayed primary fibroblasts from each species for senescence-associated β-galactosidase (SABG), which reports lysosomal hyperactivity during senescence and has served as a classic marker of senescence (44). After irradiation we detected robust signal in this assay from cells of both species, as expected; however, the proportion of SABG-positive cells in *M. spretus* fibroblast cultures was two to eight-fold lower than that of *M. musculus* cells (Figure 2). Primary fibroblasts from *M. musculus domesticus*, a close relative of *M. musculus musculus*, exhibited an intermediate SABG staining after irradiation (Supplemental Figure S1). We conclude that in the irradiated fibroblast culture model, genotypes from distinct species encode a range of lysosomal activity phenotypes, with the most avid in *M. musculus musculus*.

### The high-amplitude SASP of *M. musculus* fibroblasts is unique relative to *M. spretus*

In *M. musculus* cells, the massive lysosomal changes seen after irradiation likely result from overload of the proteostasis system during SASP production (45–47). Given that we had observed weaker effects of irradiation on lysosomal activity in fibroblasts from non-*M. musculus* species, we hypothesized that the latter would likewise exhibit a dampened-SASP phenotype. To test this, we focused on *M. musculus* and *M. spretus* as representatives of the extremes of the phylogeny. We profiled bulk RNA levels in irradiated and control primary fibroblast cultures from each species (Supplemental Table S1). In the resulting profiles, we inspected genes of the SASP immune-stimulatory program (4), and found that this gene cohort was induced more highly in *M. musculus* cells than in those of *M. spretus* after irradiation (Figure 3A). Likewise, in an unbiased search of Gene Ontology terms, we identified several suites of immune response and NF-κB signaling genes that were enriched for senescence-specific differential expression between the cultures (Figure 3B and Supplemental Tables S2 and S3). In each case, the gene groups were more strongly induced during senescence in *M. musculus* cells than in *M. spretus* cultures; among members of the latter we noted *Cxcl1* (Kim et al. 2018), *Il6* (49), *Ccls* 2,7 and 8 (50), *Mmp13* (51), and other reported SASP genes (Supplemental Figure S2). These data make clear that, at the level of mRNA, the senescence regulatory program differs markedly between fibroblasts of our focal species.

**Figure 3:**
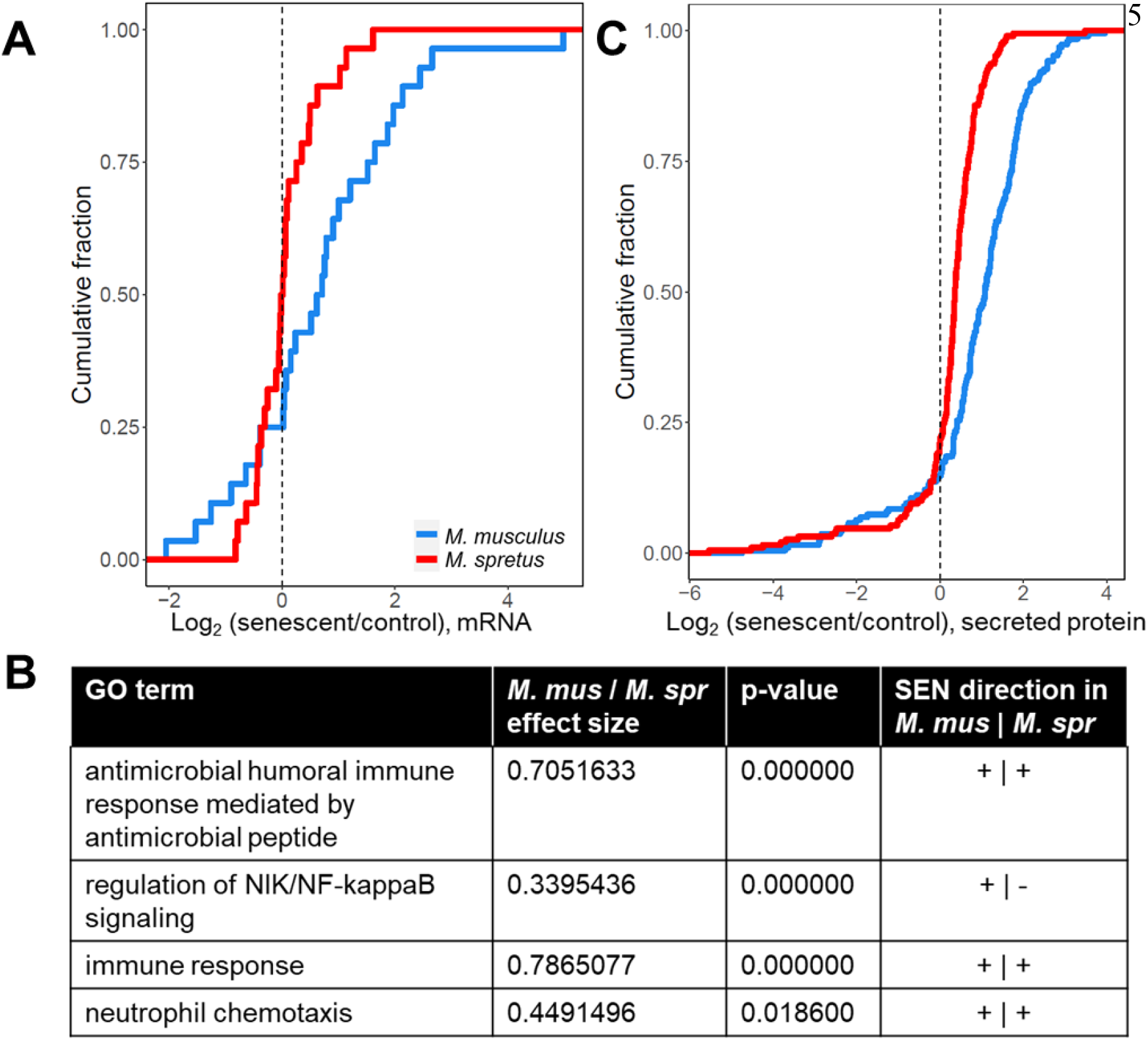
SASP is detected at higher levels in *M. musculus* cells. (A) Each trace reports a cumulative distribution of the change, in senescent primary fibroblasts of the indicated species, in mRNA levels of genes of the senescence associated secretory phenotype with senescence (Coppé et al. 2008). The *y*-axis reports the proportion of genes with the expression change on the *x*-axis, with the latter taken as an average across replicates. (B) Each row shows results from a test of the genes of the indicated Gene Ontology term for enrichment of expression change between the species during senescence, with *p*-values from a resampling-based test, corrected for multiple testing. (C) Annotations are as in (A) except that measurements were of secreted peptides, and a curated list of known SASP factors are shown (Coppé et al. 2008; Basisty et al. 2020).

We hypothesized that much of the mRNA expression divergence between *M. musculus* and *M. spretus* cells during senescence would result in differential protein abundance. In pursuing this notion, we focused on proteins secreted into the medium by senescent cells, owing to the physiological importance of the SASP (50). We collected conditioned media from senescent and control cultures of primary fibroblasts of each genotype, and we used it as input into unbiased mass spectrometry to quantify protein abundance (Supplemental Table S4). Focusing on proteins with a significant senescence-specific divergence in secretion between cells of the two species in this data source, we found higher levels overall in the medium of irradiated *M. musculus* cells relative to that of *M. spretus* (Figure 3C). This trend was borne out for a broad representation of SASP factors, including CCL chemokines, matrix metalloproteases, and serpins (Supplemental Figure S3). Together, our omics profiles reveal striking quantitative differences in SASP levels between *M. spretus* fibroblasts and those of *M. musculus*, with higher mRNA expression and protein secretion in the latter.

### A genomic screen for senescence transcription factors using cis-regulatory sequence variations

Having established divergence between *M. musculus* and *M. spretus* primary fibroblasts in senescence mRNA and protein secretion (Figure 3), we reasoned that such differences could be harnessed in an *in silico* screen for senescence regulators. We designed an analysis focused on gene regulation—in particular, on variation between the species at the binding sites of transcription factors (Figure 4A). We expected that, at some genes, *cis*-regulatory elements encoded in the *M. musculus* genome would drive expression during senescence differently than those in the *M. spretus* genome. We reasoned that if *cis*-acting variation effects were enriched among the loci bound by a given transcription factor across the genome, the signal could be interpreted as a signpost for the factor’s activity during senescence (Figure 4A). In this way, all *cis*-regulatory variants between the species that manifested in cultured primary fibroblasts, whether of large or small effect size, and regardless of their potential for phenotypic impact, could contribute to the search for transcription factors relevant for senescence.

**Figure 4:**
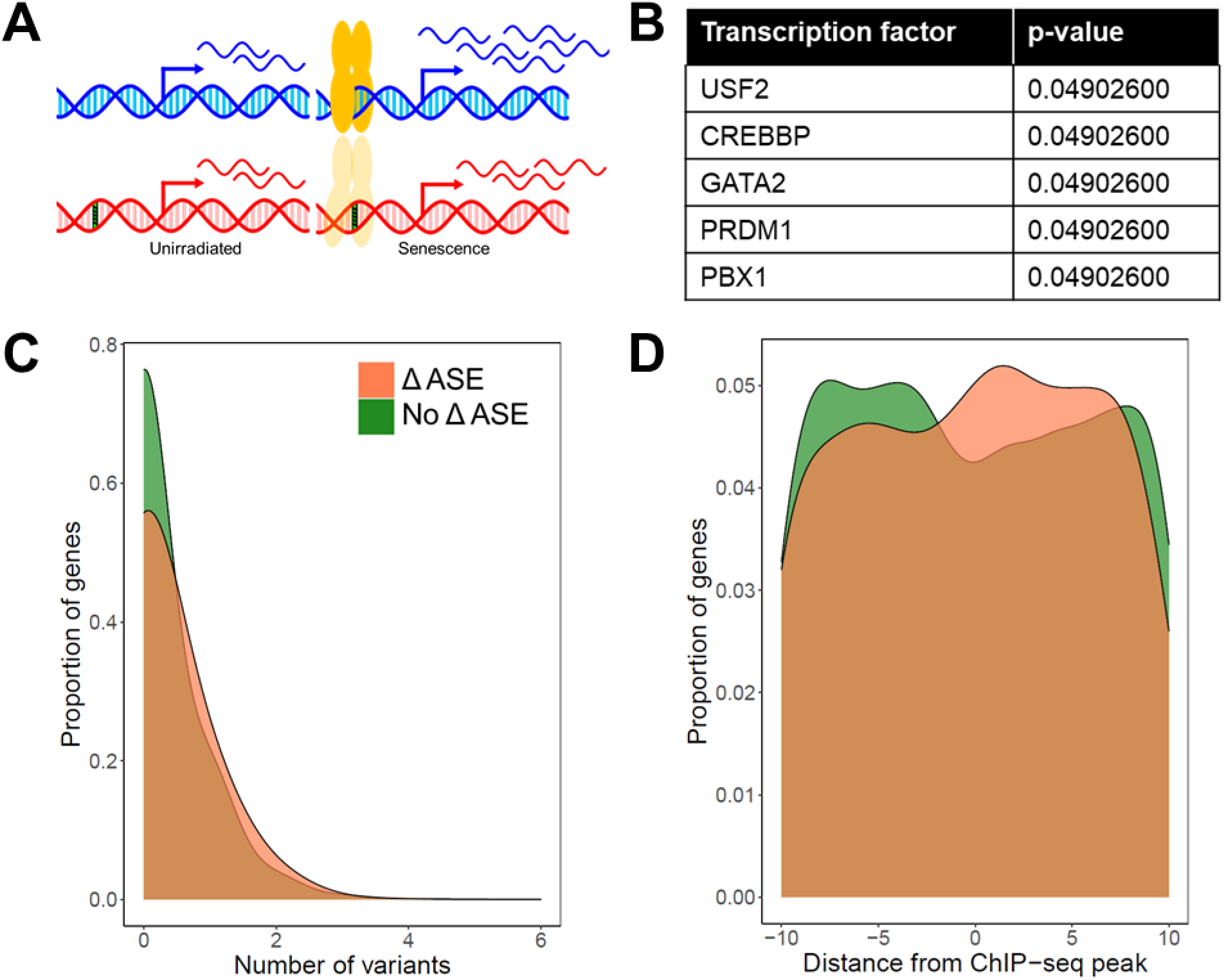
USF2 emerges as a senescence regulator candidate from a natural variation-based transcription factor screen. (A) *M. musculus* (blue) and *M. spretus* (red) alleles of a gene are expressed differently in interspecific F1 hybrid cells in a senescence-dependent manner, as a product of a sequence variant (green striped) in the binding site for a transcription factor (yellow). (B) Each row reports the multiple testing-corrected *p*-value from a Fisher’s Exact Test of target genes of the indicated transcription factor, quantifying association between species differences in experimentally determined binding sites (37) and allele-specific expression in primary cells of the *M. musculus* x *M. spretus* F1 hybrid background before and after senescence induction. Results for all tested factors are listed in Supplemental Table 5. (C) Shown are the input data for the Fisher’s Exact Test in (B) for USF2. For each trace, the *x*-axis reports the number of sequence variants between *M. musculus* and *M. spretus* in a given USF2 binding site, and the *y*-axis reports the proportion of all USF2 target genes bearing the number of variants on the *x*, as a kernel density estimate. Colors denote the presence or absence of senescence-dependent differential allele-specific expression (Δ ASE). (D) Data are as in (C) except that the *x*-axis reports the distance of the variant from the center of the USF2 binding site.

As a resource for this approach, we mated PWK *M. musculus* and STF *M. spretus* to yield F1 hybrid animals, from which we derived primary tail fibroblasts for culture and irradiation. These cells, when irradiated, exhibited a flattened morphology reflecting entry into senescence; senescence-associated β-galactosidase activity was of a magnitude between those of purebred *M. musculus* and *M. spretus* fibroblasts upon irradiation (Supplemental Figure S4). We subjected senescent and control F1 hybrid fibroblasts to RNA-seq profiling, and we used the results to quantify levels of transcripts derived from the *M. musculus* and *M. spretus* alleles of each gene in each condition (Supplemental Table S1). At a given gene, any difference between allele-specific expression in an F1 hybrid can be attributed to variants inherited from the parent species that perturb gene regulation in *cis* at the locus, because *trans*-acting factors impinge to the same extent on both alleles (52). Analyzing the response to senescence induction for a given gene, we found that the allele-specific expression difference between the alleles in the F1 hybrid was a partial predictor of the expression divergence between the *M. musculus* and *M. spretus* purebreds, in our primary cell system (Supplemental Figure S5). The latter trend reflects the joint contributions of *cis*- and *trans*-acting variants to total expression divergence between the species, as expected (53). Separately, to survey overall regulatory programs in F1 hybrid primary fibroblasts, we formulated the expression level of a given gene in a given condition as the sum of the measured levels of the *M. musculus* and *M. spretus* alleles. In this analysis, focusing on SASP genes as we had done for the purebreds (Figure 3A), we found that the expression program of senescent F1 hybrid cells was, for most components, intermediate between the low levels seen in *M. spretus* cells and the high levels in *M. musculus* (Supplemental Figure S6). These data indicate that *M. musculus* x *M. spretus* F1 hybrid fibroblasts do not exhibit heterosis with respect to senescence-associated genes, and do manifest extensive, senescence-dependent *cis*-regulatory variation.

We next used the expression measurements from *M. musculus* x *M. spretus* F1 hybrid fibroblasts as input into our *in silico* screen to identify senescence-dependent transcription factor activity. For a given transcription factor, we collated binding sites detected by chromatin immunoprecipitation upstream of genes across a panel of tissues (37). At each site, we tabulated the presence or absence of DNA sequence variants in the respective genomes of *M. musculus* and *M. spretus*. We then tested whether, across the genome, genes with these binding site variants were enriched for senescence-associated expression differences between the two alleles in the F1 hybrid. This test had the capacity for high power to detect even subtle contributions from transcription factors if they had deep binding-site coverage in the input data; five factors attained genome-wide significance (Figure 4B and Supplemental Table S5). Among them, PBX1 (54) and CREBBP (55,56) had been implicated in cellular senescence in the previous literature, providing a first line of evidence for the strength of our approach to identify signatures of condition-dependent transcription factor function. The top-scoring transcription factor in our screen, a basic-helix-loop-helix leucine-zipper protein called upstream stimulatory factor 2 (USF2), had not been experimentally characterized in stress response or senescence. However, classic studies had established USF2 as a regulator of the cell cycle and tumor suppression (57–60). More recently, USF2 was shown to control cytokine release in immune cells (61). Considering these known functions, and bioinformatic analyses suggesting a link between USF2 and senescence programs (21), we chose USF2 for in-depth validation. In detailed genomic tests, single variants between *M. musculus* and *M. spretus* at USF2 binding sites drove most of the relationship with allele-specific expression in hybrid senescent cells (Figure 4C). These variants were over-represented at positions central to, and slightly downstream of, experimentally determined peaks for USF2 (Figure 4D), highlighting the likely importance of this region in USF2’s mechanisms of binding and regulation.

### USF2 modulates cell proliferation and the acute DNA damage response

Our question at this point was whether and how USF2 regulated senescence programs. As such, we shifted our focus from natural genetic variation to controlled, laboratory-induced genetic perturbations in a single genetic background. We designed two short hairpin RNAs (shRNAs) targeting *Usf2*, each in a lentiviral vector under the U6 promoter. Expression measurements upon transformation of PWK *M. musculus* primary tail fibroblasts confirmed 2.5 and 3-fold knockdown of *Usf2* expression, respectively, from these shRNAs (Supplemental Figure S7). In these otherwise untreated cells subject to knockdown, uptake of the nucleotide analog EdU, a marker of DNA synthesis, was reduced by 40% (Supplemental Figure S8), consistent with studies of USF2 in growth of resting cells in other tissues and contexts (27,57– 60).

We now set out to use our knockdown approach to define the role of USF2 in the acute DNA damage response and senescence. For this purpose, we infected cells with *Usf2* shRNAs, cultured them under standard conditions, and then induced senescence by irradiation. DNA damage signaling, an inducer of cellular senescence (2,3), drops sharply in intensity within eight hours and then more gradually over several day after irradiation (62), culminating in a lower persistent signal (43,63–65). We first focused on the early phase of this process (six hours after irradiation) in cultures of primary fibroblasts expressing *Usf2* shRNAs or scrambled shRNA controls (Figure 5A). Transcriptional profiling followed by Gene Ontology analyses identified gene groups enriched for expression changes dependent on condition and USF2 (Figure 5B and Supplemental Tables S6 and S7). Most salient was a trend of pervasive repression transcriptome-wide six hours after irradiation, which was detected in control cells as expected (66,67), and was blunted in cells with *Usf2* knocked down. The latter effect was particularly enriched in the transcriptional machinery, repressors of apoptosis, and several cell proliferation regulators (Figure 5B-D and Supplemental Figure S9). DNA repair genes, a likely target for changes upon irradiation, are regulated primarily at the post-transcriptional level (68,69), and we did not detect effects of *Usf2* knockdown on their transcripts (Supplemental Figure S10).

**Figure 5:**
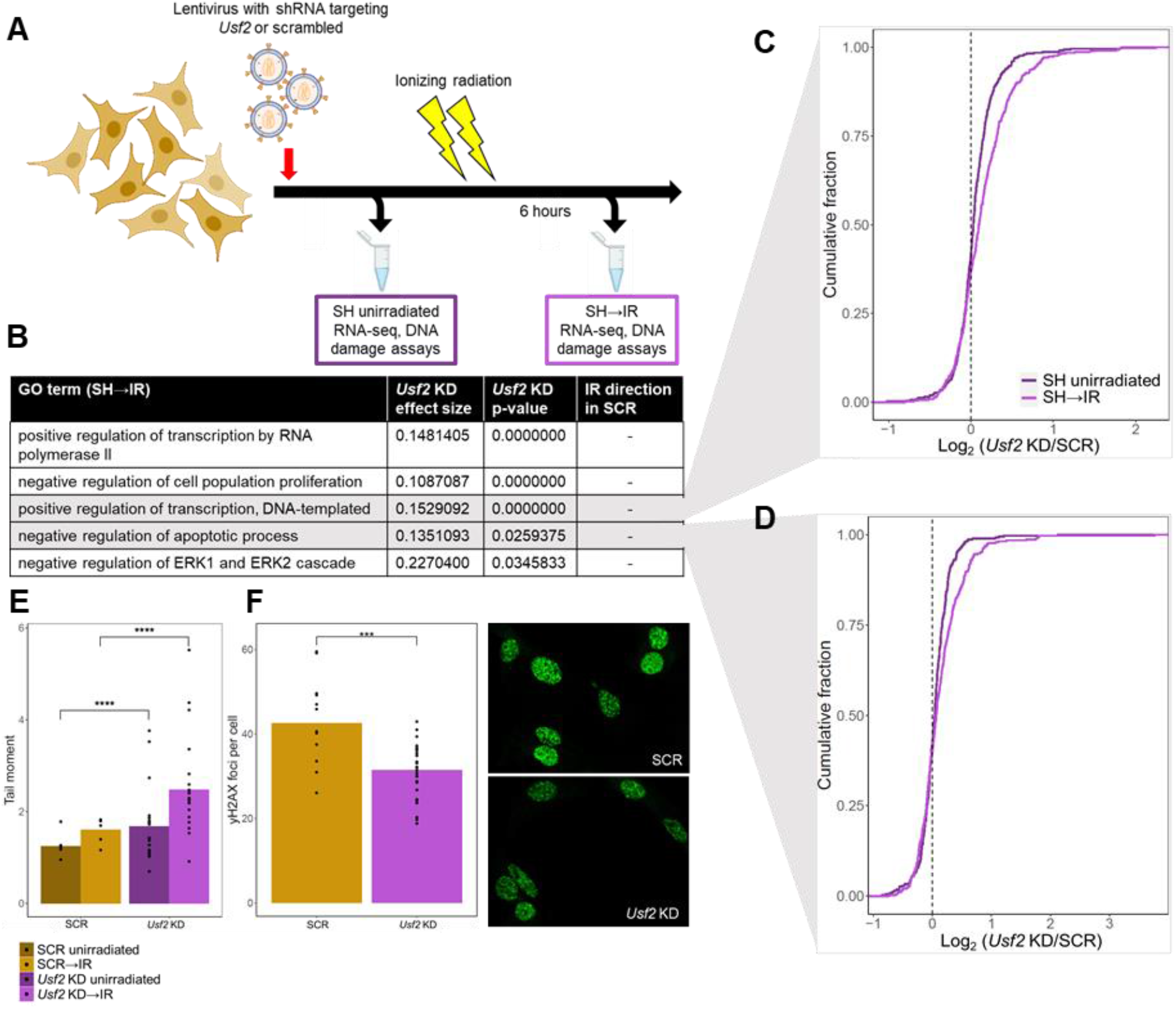
*Usf2* depletion results in more DNA damage but a muted DNA damage response following irradiation. (A) *M. musculus* primary fibroblasts were infected with a lentivirus encoding a short hairpin RNA (shRNA, SH) targeting *Usf2* or a scrambled control (SCR), and analyzed before (SH unirradiated) or six hours after (SH→IR) treatment with ionizing radiation. (B) In a given row, the second column reports the average, across genes of the indicated Gene Ontology term, of the log_2_ of the ratio of expression between *Usf2* knockdown (KD) and SCR-treated cells, six hours after irradiation. The third column reports significance in a resampling-based test for enrichment of directional differential expression between *Usf2* KD and SCR-treated cells in the respective term, corrected for multiple testing. The fourth column reports the direction of the change in expression six hours after irradiation in SCR-treated cells. (C) Each trace reports a cumulative distribution of the log_2_ of the ratio of expression between *Usf2* KD or SCR-treated cells in genes annotated in the positive regulation of transcription, before or six hours after irradiation treatment as indicated. The *y*-axis reports the proportion of genes with the expression change on the *x*-axis. (D) Data are as in (C), except that genes involved in apoptosis were analyzed. (E) Each column reports tail moments detected in a comet assay on primary fibroblasts harboring the indicated shRNAs, before or six hours after irradiation. In a given column, points report biological and technical replicates and the bar height reports their average (SCR *n* = 5, *Usf2* KD *n* = 20). ****, *p* < 0.0001, one-tailed Wilcoxon test. (F) Left, each column reports number of γH2AX foci per cell detected in primary fibroblasts harboring the indicated shRNAs six hours after irradiation. Data are displayed as in (E) (SCR *n* = 12, *Usf2* KD *n* = 28). ***, *p* < 0.001, one-tailed Wilcoxon test. Right, representative images of the indicated cultures.

We hypothesized that *Usf2* knockdown during the acute DNA damage response would also have cell-physiological effects. Assays of EdU incorporation to report on DNA synthesis showed effects of *Usf2* knockdown after irradiation to the same degree as in resting cultures (Supplemental Figure S7). To focus on phenotypes more proximal to DNA damage, we used the neutral comet assay (39) to measure DNA double-stranded breaks on a per-cell basis. In this setup, *Usf2* depletion increased comet tail moments by 50% six hours after irradiation, with an effect that was similar, though of smaller magnitude, in resting cell controls (Figure 5E). Next, we tracked foci of phosphorylated histone H2AX (γH2AX) in fibroblasts as a marker of chromatin decondensation, preceding the repair of DNA double-stranded breaks (70). Cells harboring *Usf2* shRNAs exhibited 30% fewer γH2AX foci than cells of the control genotype, six hours after irradiation (Figure 5F). These data establish a role for USF2 in the response to irradiation, with knockdown of this factor compromising cells’ ability to mount the classical transcriptional program under this stress, and to carry out DNA damage repair.

### USF2 tunes the commitment to senescence

Having knocked down *Usf2* in PWK primary fibroblasts and irradiated them to study the acute DNA damage response, we now allowed the irradiated cultures to enter senescence (Figure 6A). We referred to this as a “knockdown-then-irradiate” experimental design (SH -> SEN in Figure 6). Growth arrest and flattened morphology were indistinguishable between these cells and controls harboring scrambled shRNAs (see Figure 6C), indicating that wild-type levels of *Usf2* were not required at the point of irradiation to establish senescence *per se*. To investigate quantitative characteristics of these senescent cultures, we subjected them to expression profiling and Gene Ontology enrichment analyses (Figure 6B and Supplemental Tables S6 and S8). Among top-scoring gene groups, the most dramatic effects were in those that dropped in expression in senescent cultures of the control genotype, which as expected (17,71) included cell cycle and DNA repair pathways (Figure 6B and Supplemental Table S8). Intriguingly, mRNA levels of the latter were even lower in senescent cells that had been irradiated after *Usf2* knockdown, showing a reduction of ~20% on average (Figure 6B). Among the genes of this cohort, some of which declined in expression by >5-fold with *Usf2* knockdown in senescence, we noted cell cycle regulators (*Ccna2*, *Cdc20*, *Cdk1)*, kinesin components (*Kif2c, Knl1*), the DNA polymerase *Pole*, and the DNA damage checkpoint ubiquitin ligase *Uhrf1* (Supplemental Figure S11A). We conclude that genes of the cell cycle and DNA repair machinery are detectable at a low but non-zero expression level in wild-type senescent cells, and that these pathways are subject to further reduction when *Usf2* is limiting.

**Figure 6:**
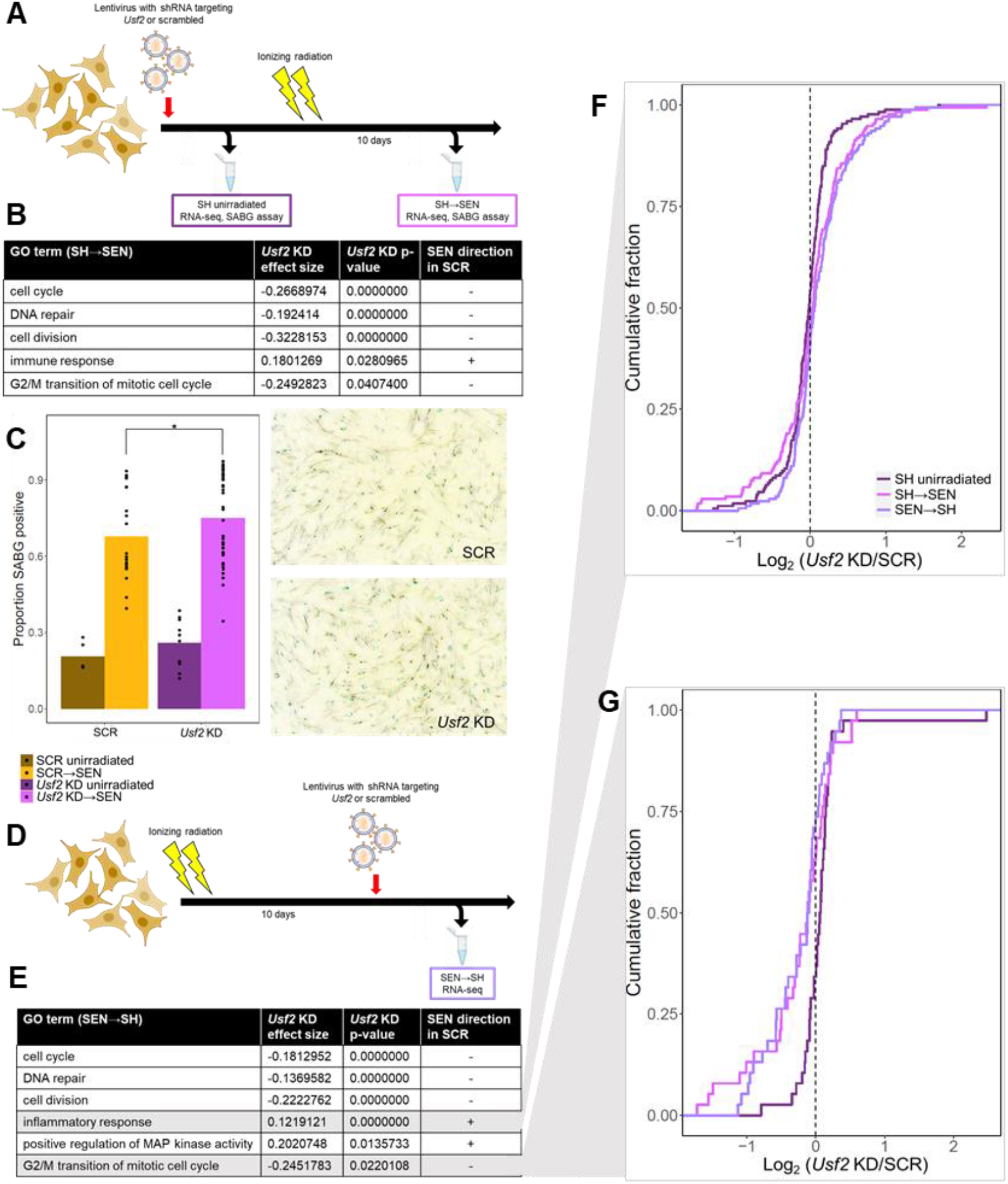
*Usf2* knockdown results in an enhanced senescence profile. (A) *M. musculus* primary fibroblasts were infected with a lentivirus encoding an shRNA (SH) targeting *Usf2* or a scrambled control, and analyzed before (SH unirradiated) or after (SH→SEN) treatment with ionizing radiation (IR) to induce senescence (SEN). (B) Data are as in Figure 4B except that cells were analyzed 10 days after irradiation. (C) Left, each column reports the proportion of senescence associated β-galactosidase (SABG)-positive cells treated with the indicated shRNAs in resting culture (unirradiated) or 7 days after irradiation (SEN) as in (A). In a given column, points report biological and technical replicates and the bar height reports their average (SCR unirradiated *n* = 5, SCR SEN *n* = 21, *Usf2* KD unirradiated *n* = 10, *Usf2* KD SEN *n* = 42). *, *p* < 0.05, one-tailed Wilcoxon test. Right, representative images of the indicated cultures. (D) *M. musculus* primary fibroblasts were irradiated, incubated for 10 days to senesce, then infected with shRNAs and analyzed after 10 additional days. (E) Data are as in (B) except that cells from the scheme in (D) were analyzed. (F) Each trace reports a cumulative distribution of the log_2_ of the ratio of expression in *Usf2* knockdown (KD) and scrambled control (SCR)-treated cells, in genes annotated in the inflammatory response, when shRNAs were administered to a resting culture (SH unirradiated), to resting cells followed by irradiation as in (A) (SH→SEN), or after irradiation and senescence establishment as in (C) (SEN→SH). The *y*-axis reports the proportion of genes with the expression change on the *x*-axis, with the latter taken as an average across replicates. (G) Data are as in (F), except that genes involved in G2/M transition of mitotic cell cycle were analyzed.

Likewise, *Usf2* knockdown also affected inflammation and immune-recruitment factors (Figure 6B and Supplemental Table S8). Cells of the control genotype induced these pathways during senescence, as expected (2,3,9,49,72,73); in cells reaching senescence after *Usf2* knockdown, induction of inflammatory factors was amplified by ~10% on average (Figure 6B and Supplemental Figure S11B). Similarly, in assays of the β-galactosidase senescence marker, we observed a 10% increase in cells subject to *Usf2* knockdown and senescence treatment (Figure 6C). Together, our profiling data establish that irradiation of primary fibroblasts with reduced *Usf2* expression leads to a quantitatively perturbed, exaggerated senescent state, with reduced expression of proliferation and DNA repair pathways, and elevated pro-inflammatory gene expression and β-galactosidase activity.

We reasoned that the changes in senescence we had seen upon irradiation of *Usf2*-depleted cells could constitute, in part, effects from increased DNA damage in the knockdown genotype (Figure 5). To pursue the role of USF2 in senescence more directly, we used a distinct experimental paradigm: we irradiated wild-type cells and incubated them for 10 days to allow establishment of senescence, and we then expressed shRNAs targeting *Usf2*, followed by 10 additional days of incubation (Figure 6D). We referred to this as an “irradiate-then-knockdown” experiment (SEN -> SH in Figure 6). We first inspected control cells for this paradigm, harboring a scrambled shRNA. Here RNA-seq revealed expression changes for many genes from the resting state through irradiation and early and late senescence (Supplemental Tables S6 and S9, and Supplemental Figure S12), attesting to the dynamics of senescence as expected (17,71). We next carried out RNA-seq and GO term enrichment analysis of cultures subject to *Usf2* knockdown during late senescence (Figures 6D-G and Supplemental Tables S6 and S10), for comparison to our “knockdown-then-irradiate” strategy (Figure 6A). On average, cell cycle and DNA repair genes, repressed in control cells during senescence, were expressed at even lower levels when *Usf2* was knocked down midway through the senescence time course; this was analogous to our findings upon early knockdown of *Usf2* (compare lavender to magenta in Figure 6G and Supplemental Figure S11A). Likewise, inflammatory response factors, induced during senescence in the control setting, were more highly expressed in the “irradiate-then-knockdown” approach, consistent with our findings from the early-knockdown design (compare lavender to magenta in Figure 6F and Supplemental Figure S11B). We noted only a handful of genes for which early *Usf2* knockdown effects were not recapitulated in our paradigm of knockdown after senescence entry (*e.g. Adamts1* and *Rarres2* in Supplemental Figure S11B). Overall, our analyses establish USF2 as a senescence regulator at least in part independent of its role in the acute DNA damage response, such that in its absence, cells commit even more strongly to the senescent state.

## Discussion

Complex regulatory networks likely underlie many of the quantitative behaviors of senescent cells, including kinetics and dependence on cell type and inducer (15,17,26,74,75). Exactly how these nuances are encoded remains poorly understood. In this study, we pioneered the use of interspecies genetic divergence to screen for components of the senescence regulatory machinery. This strategy complements previous studies of transcription factor binding sites in genes of the senescence program in a single genetic background (17,21). Our approach harnesses the correlation between interspecies variation in sequence and expression levels, as an additional line of evidence for senescence-specific regulatory functions by a given factor. This paradigm parallels similar tools previously used to dissect divergence in expression (76,77) and transcription factor binding (78–80) in other contexts. Broadly speaking, these methods are not highly powered for pathways under strong evolutionary constraint, which, by definition, will not vary enough among species to yield the raw observations that would go into a screening pipeline. Rather, we expect the natural variation-based approach to work best for discovering less-constrained modifiers, many of which may confer layers of quantitative regulation onto a master regulatory pathway.

We focused our experimental validation on one such modifier, the transcription factor USF2. By tracing USF2’s function in proliferation and genome-wide expression in untreated cells, we extended conclusions from studies of USF2 in tumor suppression and cell cycle regulation (57– 59,61), apoptosis (81) and ERK1/2 signaling (82). In an acute DNA damage setting, we discovered that USF2 is required for cells to mount DNA repair and downstream DNA damage responses. And in senescence proper, we showed that USF2 acts as a repressor, such that in its absence, the senescence program—shutoff of cell proliferation and DNA repair, and induction of cytokines—is amplified. A compelling model is thus that even long after damage exposure, cells have access to expression states along a continuum of commitment to senescence, and that USF2 acts to help determine which state they occupy. If so, USF2 would take a place among a network of factors, including p53, ING, Rb (83,84), p21 (85,86), and p16 (87), that govern the choice between senescence, apoptosis, and repair and proliferation, depending on cell type (83) and the amount of damage or stress incurred (84).

As a corollary of these conclusions from expression profiling, we note that cell cycle and DNA repair genes, classically known to be repressed during senescence (17,71), did not hit a floor of expression in senescent cultures: we could detect them at even lower expression levels upon *Usf2* knockdown. Since our cultures comprise >99% arrested cells within several days of irradiation (see Supplemental Methods), the emerging picture is that the proliferation machinery is maintained at non-zero levels even in such a population. Any ability of these gene products to reattain activity could be of particular interest as a potential mediator of the return to proliferation seen among senescent cells in certain scenarios (88,89).

Our work leaves open the mechanisms by which USF2 exerts its effects in the DNA damage response and cellular senescence. It is tempting to speculate that USF2 ultimately works in these processes in concert with its better-studied family member, USF1. Indeed, USF1 has been implicated in DNA repair (90), inflammation (91,92), immune responses (93), and p53-mediated cell cycle arrest (94) in contexts other than senescence. In addition, given that USF2 has been implicated in the TGFβ-p53 axis in apoptosis (81) and fibrosis (95), the latter pathway could mediate some part of the USF2 effects we have seen. Furthermore, regulatory network reconstruction (41) suggests that USF2 acts upstream of several other transcription factors (KLF3, GLI3, NFIL3) with direct targets in DNA repair, DNA damage response, and senescence pathways (Supplemental Table S11).

Alongside our use of *cis*-regulatory variation between mouse species as a screening tool for senescence genes, we also characterized overall patterns of divergence between *M. spretus* senescent cells and those of *M. musculus*. Given that the former exhibited lower levels of SASP mRNAs and proteins, we suggest that the rheostat of the senescence response is at a higher set point in this species, such that at a given level of stress (*e.g*. the irradiation we study here), cells of this species synthesize and secrete less of the SASP. Under this model, the decision set point for commitment to senescence by irradiated cells is similar across species, and, considered at any given time after damage exposure, it is the amplitude of the SASP that has been tuned by evolution. Such an idea would have precedent in the gradual ramp-up of senescence expression in *M. musculus* cells (17): plausibly, *M. spretus* could be hard-wired for slower kinetics of this progression, in the fibroblasts we study here. *M. spretus* cells could also simply cap the amplitude of their SASP, limiting the immune recruitment function of senescent cells at any timepoint.

We further hypothesize that the dampened SASP might be a proximal cause for the lower senescence-associated β-galactosidase activity we have seen in *M. spretus* cells. Such a link would follow from current models of the senescent state in which production and secretion of SASP components (96) leads to proteotoxic stress from insoluble aggregates (45,46), an increase in the number and size of lysosomes (97), and enhanced β-galactosidase activity (98). Plausibly, any of the phenotypes we study here in cell culture could have consequences *in vivo*, with potential links to the stress- and pathogen-resistance phenotypes characterized in *M. spretus* (99–103).

The low-amplitude senescence program we have seen in *M. spretus* provides an intriguing contrast to the trend for fibroblasts from naked mole rat in culture to avoid both senescence and apoptosis altogether, after irradiation (104). Instead, a given naked mole rat cell can often resolve DNA damage sufficiently to re-enter the cell cycle, to a degree several-fold beyond that seen in *M. musculus*. Likewise, the beaver allele of the DNA damage factor SIRT6 confers a similar effect in a heterologous system (105). These represent evolutionary innovations in other rodents distinct from the quantitative tuning of senescence expression we have traced in *Mus*. The emerging picture is one in which no single irradiation response mechanism manifests in all species, even in the simplest cell culture systems. Indeed, against the backdrop of the classic literature on *M. musculus* senescence (49,106), many other irradiation response behaviors may remain to be discovered in additional non-model species. Human cells exhibit an avid senescence response, on par with that of *M. musculus* (49). As such, the programs nature has invented in other lineages may hold promise in the search for therapeutics that would tamp down the pro-aging effects of senescence in a clinical context.

## Ethics declarations

*Ethics approval and consent to participate*: All methods used in animal husbandry and sample collection were approved under Montana Institutional Animal Care and Use Committee protocol number 062-1JGDBS-120418 and reported in accordance with the ARRIVE guidelines (https://arriveguidelines.org). All other methods described were performed in accordance with the rules and regulations of the corresponding institutions.

## Consent for publication

Not applicable.

## Availability of data and materials

The datasets supporting the conclusions of this article are available in the NCBI Gene Expression Omnibus (GEO; https://www.ncbi.nlm.nih.gov/geo/) under accession number GSE201217. Raw data and complete MS data sets have been uploaded to the Mass Spectrometry Interactive Virtual Environment (MassIVE) repository, developed by the Center for Computational Mass Spectrometry at the University of California San Diego, and can be downloaded using the following link: http://massive.ucsd.edu/ProteoSAFe/status.jsp?task=5e7aa6a2b31f4dcfafa679e9d3f3d9c3 (MassIVE ID number: MSV000089246; ProteomeXchange ID: PXD033182), with username: MSV000089246_reviewer, and password: winter.

## Acknowledgements

The authors thank Mary West for support of cell culture resources, Melissa Chao for help with data collection, Michael Nachman for *M. domesticus* animal material, and Herbert Kasler, Marius Walter, and Eric Verdin for helpful discussions and materials.

## Author contributions

RBB and TK conceived of the idea of the study; TK carried out fibroblast harvest and culture, RNA-seq, knockdown, and cell-biological assays and all data analysis. BS and CDK carried out proteomics profiling. ECKK, ECM, and JMG carried out all mouse mating and husbandry. TK, JC, and RBB wrote the manuscript with input from all authors.

