## Supplemental figures for "A natural variation-based screen in mouse cells reveals USF2 as a regulator of the DNA damage response and cellular senescence"

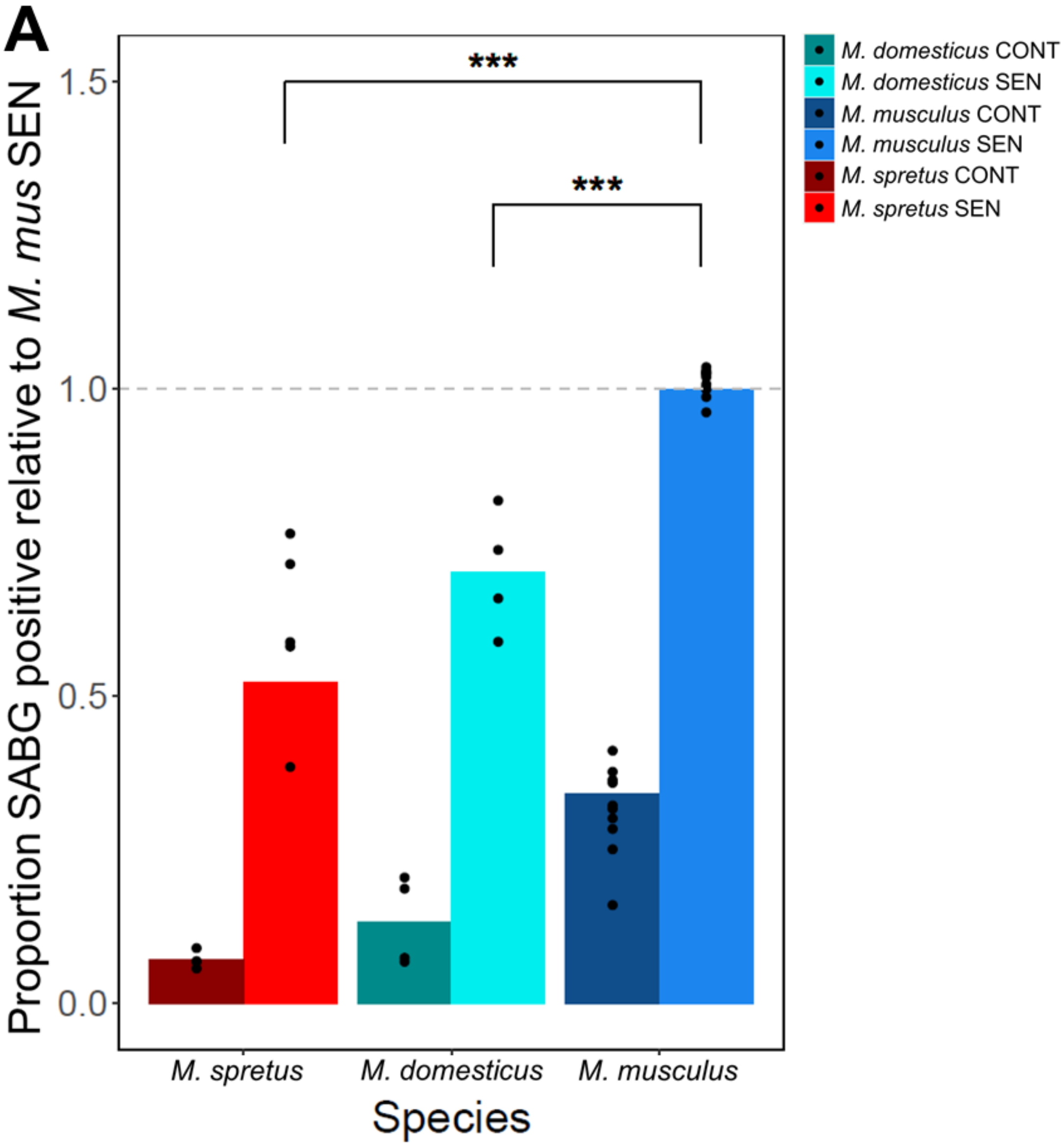


**Supplemental Figure S1: *M. musculus* and *M. spretus* cells represent the extremes in the natural variation of senescence-associated β-galactosidase across *Mus*.** (A) Each bar reports the average proportion of senescence-associated β-galactosidase (SABG) positive cells set relative to the value in senescent (SEN) M. musculus cells, for both senescent and unirradiated controls (CONT) of each species as described on the x-axis. For a given column, points represent technical replicates (*M. musculus* $n$ = 11, *M. domesticus* $n$ = 4, *M. spretus* $n$ = 5). ***, $p < 0.001$, one-tailed Wilcoxon comparing species in senescence.

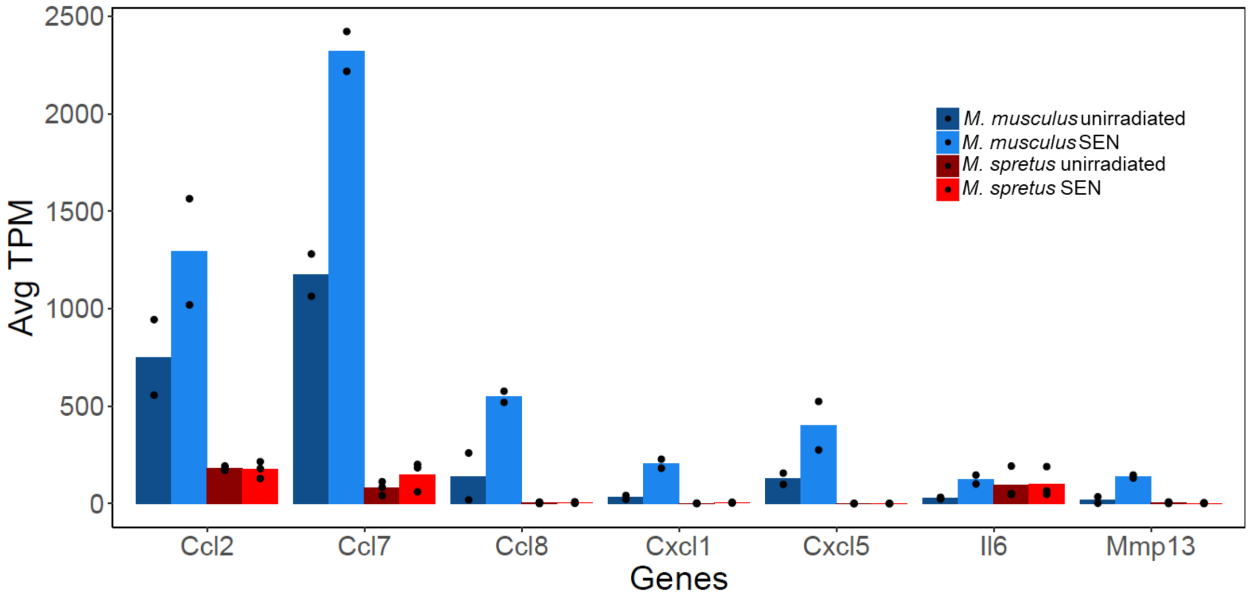


**Supplemental Figure S2: Senescent *M. musculus* primary fibroblasts display enhanced mRNA induction of genes of the senescence-associated secretory phenotype.** Each column reports expression (TPM, transcripts per million) from RNA-seq profiling of primary fibroblasts from the indicated species for the indicated genes, in control or irradiation-induced senescent (SEN) cells. In a given column, points report biological replicates and the bar height reports their average ($n$ = 3).

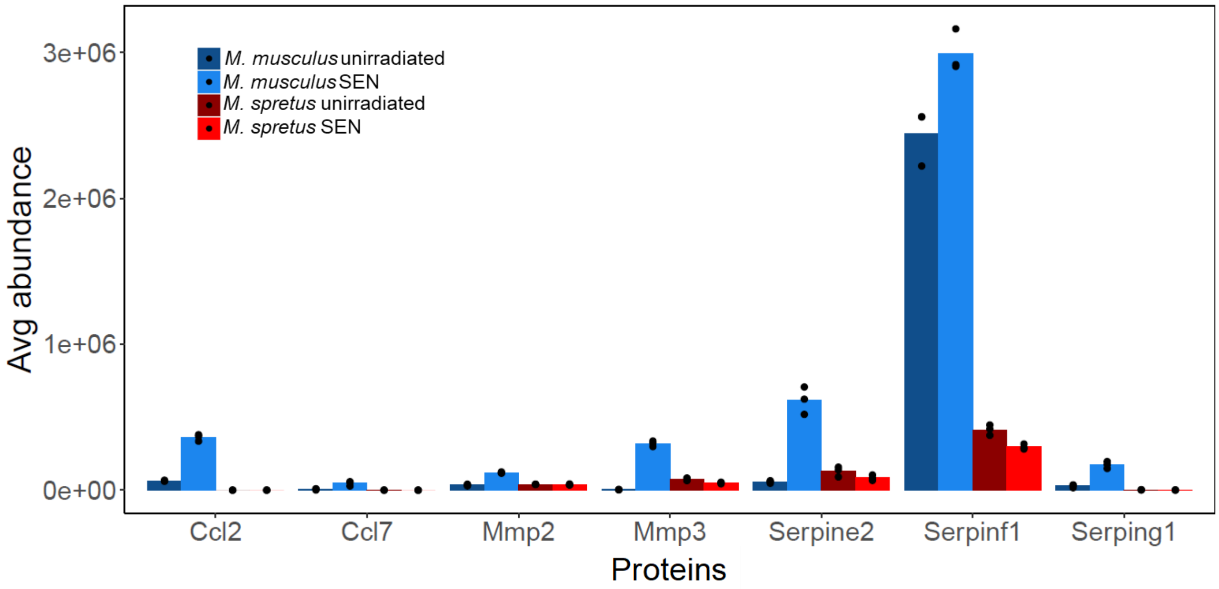


**Supplemental Figure S3: Senescent *M. musculus* primary fibroblasts display enhanced secretion of proteins of the senescence-associated secretory phenotype.** Data are as in Supplemental Figure S2 except that protein abundance from conditioned medium is shown.

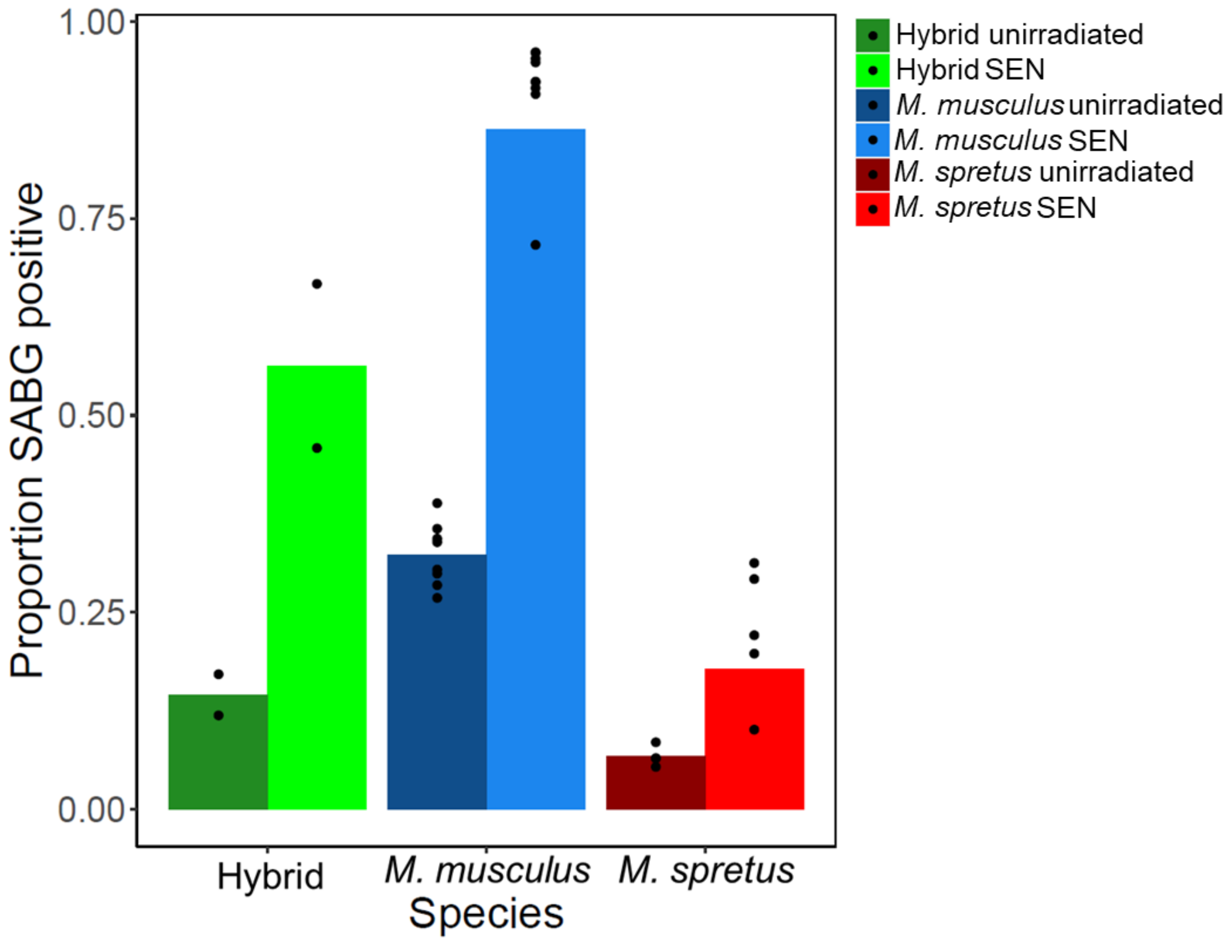


**Supplemental Figure S4: Senescent *M. musculus* x *M. spretus* F1 primary fibroblasts display intermediate activity of senescence-associated β-galactosidase (SABG).** Each column reports the proportion of SABG positive cells for the indicated genotype in control cells or in senescent cells seven days after irradiation (SEN). In a given column, points report biological and technical replicates and the bar height reports their average (*M. musculus* $n = 9$, *M. spretus* $n = 5$, F1 hybrid $n = 2$).

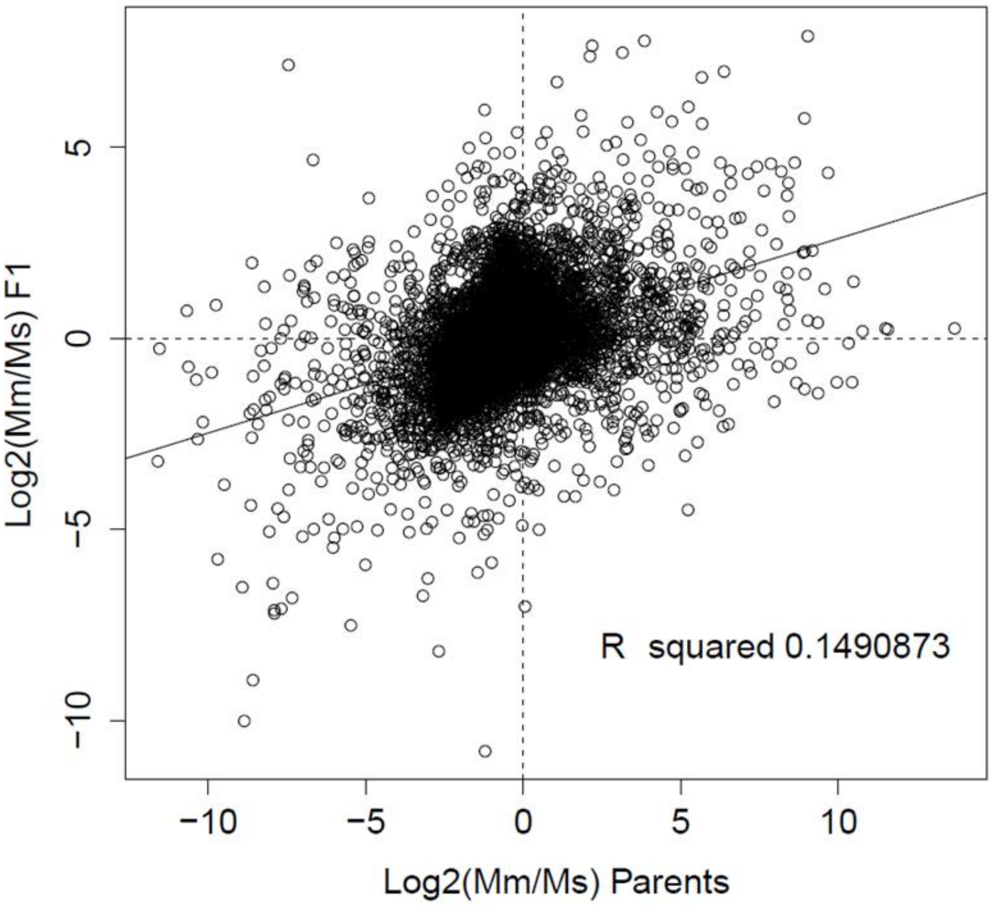


**Supplemental Figure S5: Variation in gene expression during senescence between *M. musculus* and *M. spretus* is controlled by both *cis*- and *trans*-acting elements.** Each point represents expression of one gene in primary fibroblasts induced to senesce. The *x*-axis reports the ratio of expression, as an average across biological replicates, measured in purebred *M. musculus* (Mm) and *M. spretus* (Ms) cells; the *y*-axis reports the ratio of allele-specific expression from the *M. musculus* and *M. spretus* alleles in cells from the interspecific F1 hybrid.

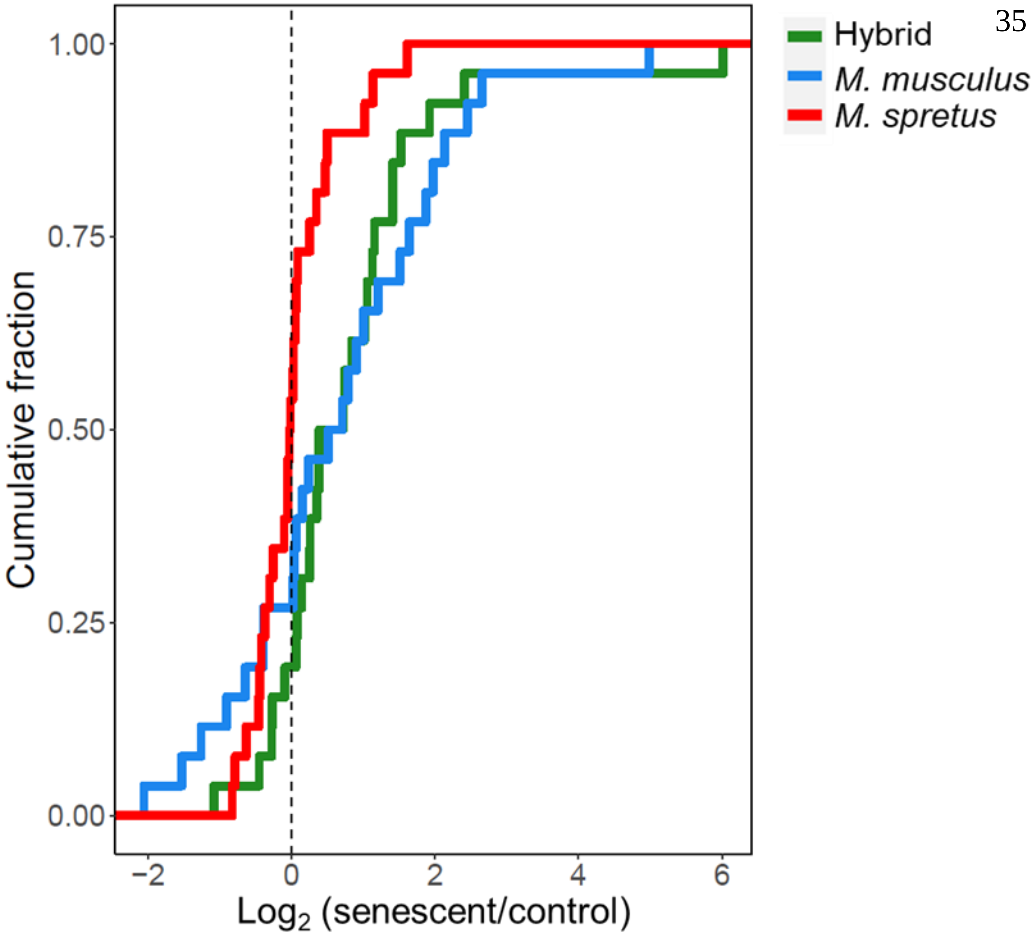


**Supplemental Figure S6: Senescent *M. musculus* x *M. spretus* F1 primary cells display intermediate induction in mRNA expression of genes of the senescence-associated secretory phenotype.** Data are as in Figure 3A of the main text except that measurements from the interspecific F1 hybrid are shown in green.

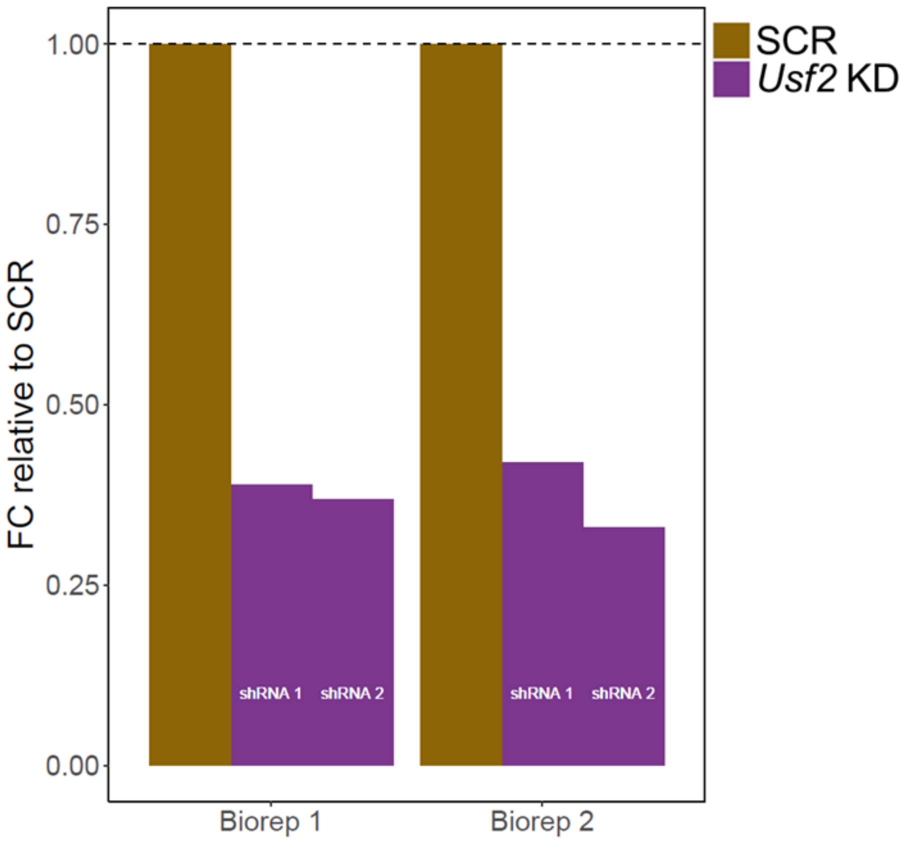


**Supplemental Figure S7: *Usf2*-targeting shRNAs knock down *Usf2* expression.** Each set of bars reports *Usf2* expression measured via qPCR in *M. musculus* primary cells, in one biological replicate. In a given replicate, each bar reports the fold change (FC) in *Usf2* expression between cells harboring *Usf2*-targeting shRNA (*Usf2* KD) and those with a scrambled shRNA (SCR), normalized with respect to the value of the latter.

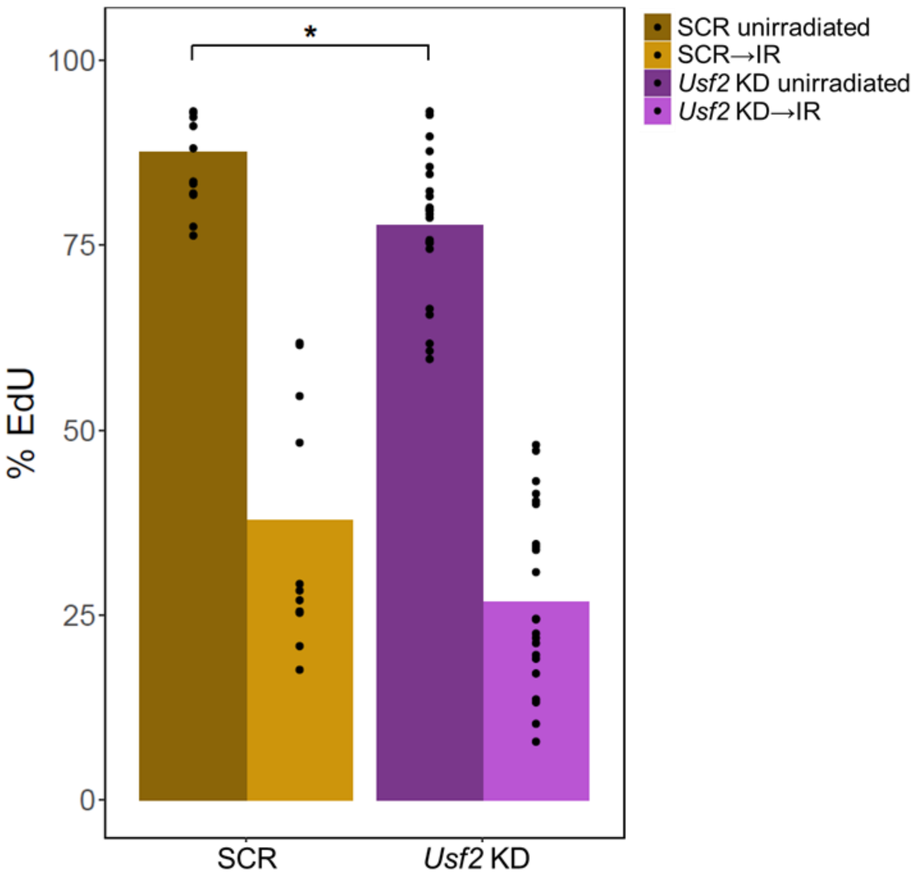


**Supplemental Figure S8: *Usf2* depletion slightly slows growth of primary *M. musculus* cells independent of irradiation.** Each column reports the percentage of EdU incorporation in primary fibroblasts harboring *Usf2* or scrambled shRNAs, before or six hours after irradiation as indicated. *, $p < 0.05$, one-tailed Wilcoxon test. In a given column, points report biological and technical replicates and the bar height reports their average (SCR $n = 11$, *Usf2* KD $n = 22$).

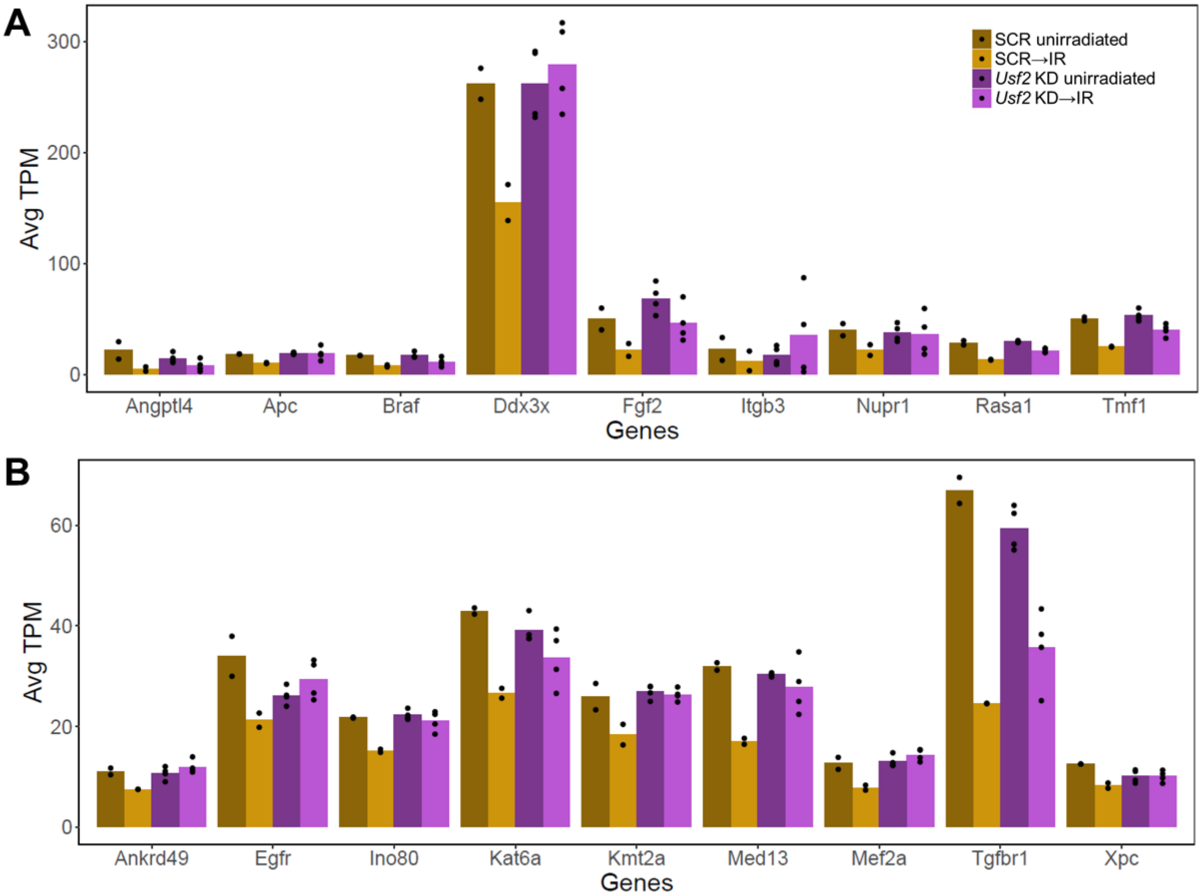


**Supplemental Figure S9: *Usf2* knockdown results in less transcriptional repression following acute DNA damage.** In a given panel, each column reports mRNA expression of the indicated gene in transcripts per million (TPM) in primary fibroblasts harboring *Usf2* or scrambled shRNAs, before or six hours after irradiation as indicated. (A) Shown are a representative subset of the genes in the Gene Ontology (GO) term “negative regulation of apoptotic process” that were repressed in cells harboring the scrambled shRNA control following acute DNA damage. (B) Shown are genes in the GO term “positive regulation of transcription, DNA-templated”. In a given column, points report biological replicates and the bar height reports their average ($n$ = 3).

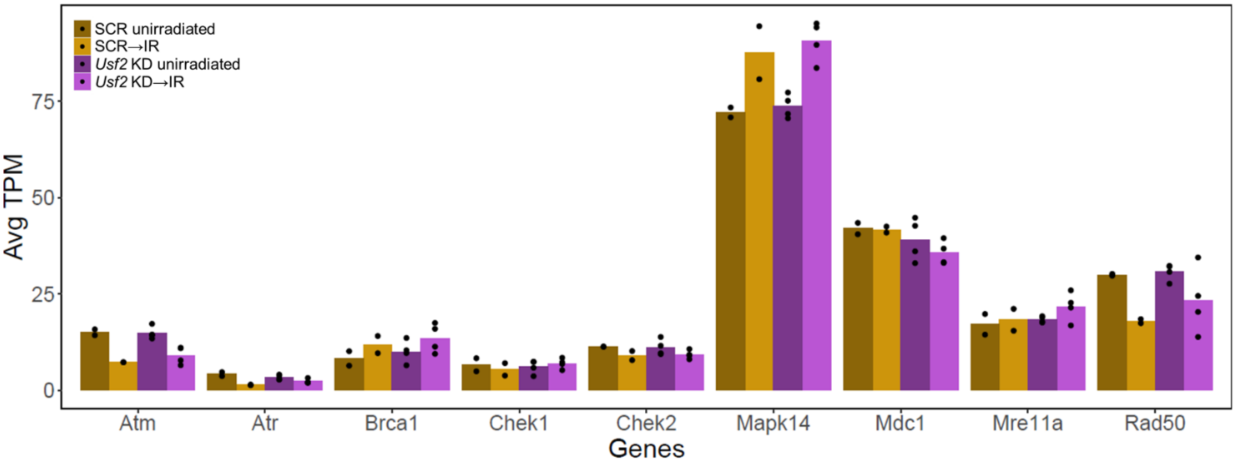


**Supplemental Figure S10: Expression of core DNA damage response genes are largely not affected by *Usf2* knockdown.** Data are as in Supplemental Figure S9 except that DNA damage response genes are shown.

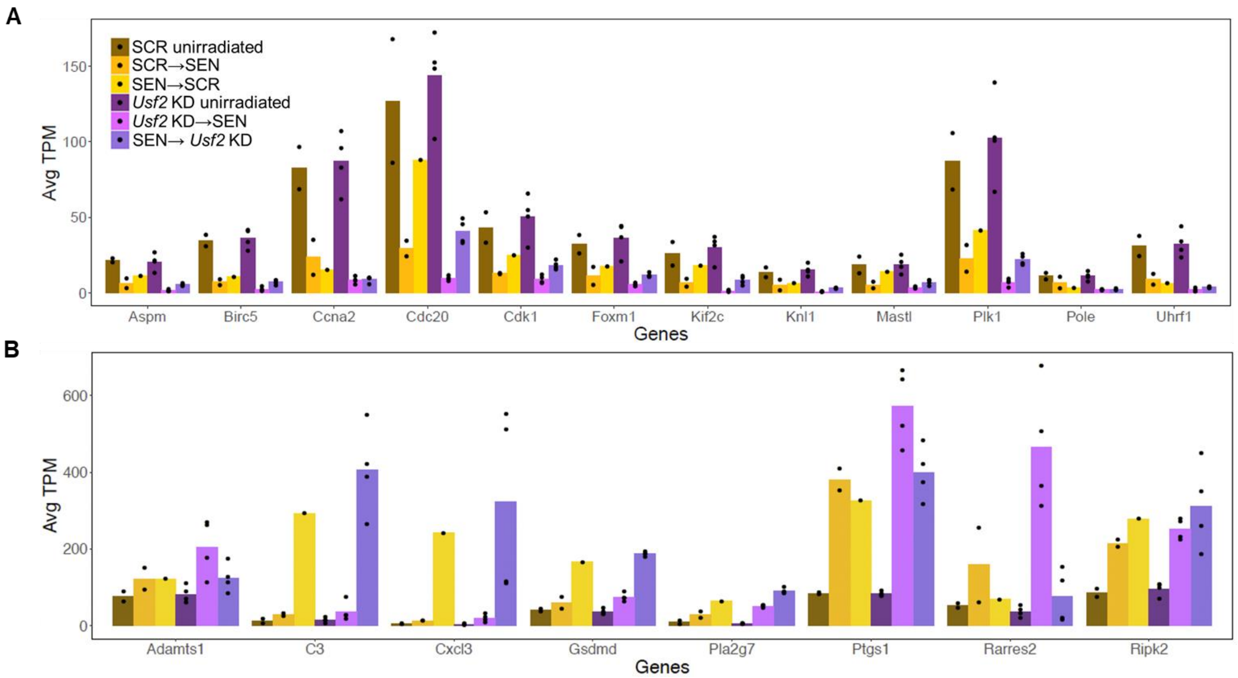


**Supplemental Figure S11: *Usf2* knockdown results in an enhanced senescent gene expression profile.** In a given panel, each column reports mRNA expression in transcripts per million (TPM) in primary fibroblasts harboring *Usf2* or scrambled shRNAs, when shRNAs were administered to a resting culture (unirradiated), to resting cells followed by irradiation (→SEN), or after irradiation and senescence establishment (→KD). (A) Genes annotated in inflammation and immune response that were upregulated during senescence in scrambled shRNA controls. (B) Genes annotated in cell cycle that were repressed with senescence in scrambled shRNA controls. In a given column, points report biological replicates and the bar height reports their average (SCR *n* = 2, *Usf2* KD *n* = 4).

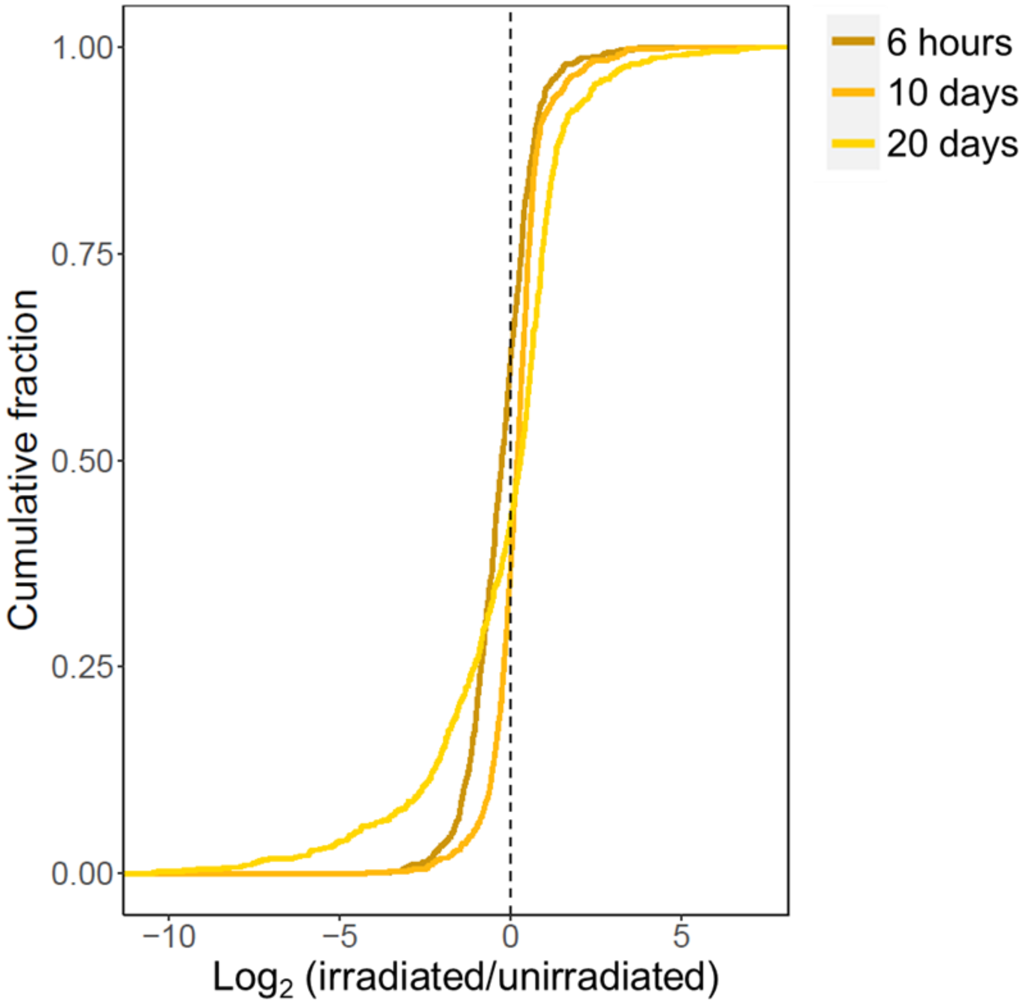


**Supplemental Figure S12: Expression profiling of control cells during a senescence timecourse reveals a dynamic expression program with senescence progression.** Each trace reports a cumulative distribution of gene expression in primary *M. musculus* fibroblasts harboring a scrambled shRNA, at the indicated timepoint after irradiation treatment relative to unirradiated controls. Included in the distribution are all genes significant in a test for significant expression change across the timepoints by multivariate ANOVA (Supplemental Table S9). The *y*-axis reports the proportion of genes with the expression change on the *x*-axis.
